# Eukaryotic tRNA ligases mediate RNA break repair

**DOI:** 10.64898/2026.05.26.727988

**Authors:** Alexander N. Wirth, Isabel S. Naarmann-de Vries, Anne M. Pinnen, Aiswarya Gopal, Anna Righetti, Kathrin Leppek, Christoph Dieterich, Jirka Peschek

## Abstract

RNA is continuously exposed to damage during physiological metabolism and stress, yet cellular responses to RNA damage remain less understood than DNA repair pathways. RNA strand breaks are particularly deleterious because they generate chemically incompatible RNA ends. Eukaryotic tRNA ligases have been implicated in RNA processing and repair, but whether they function as general RNA repair enzymes remains unresolved. Here, we show that the evolutionarily divergent tRNA ligases, human RTCB and fungal Trl1, mediate RNA break repair (RBR) targeting ribosomes and other ribonucleoprotein (RNP) complexes. Using direct RNA nanopore sequencing, we map these repair events at nucleotide resolution, demonstrating that tRNA ligases repair breaks in ribosomal RNA and restore translational activity of repaired ribosomes. We further identify repair across additional structured cellular RNAs. We show that loss of RBR activity leads to RNA fragmentation in human cells and impairs cell viability upon oxidative stress. Together, these findings uncover a broader role for eukaryotic tRNA ligases in repairing RNA breaks and maintaining transcriptome integrity.

## INTRODUCTION

RNA molecules are inherently prone to damage and cleavage during normal cellular processes and stress. Such damage can create acute biological conflicts, with severe consequences for cellular physiology^1–3^. Preserving RNA integrity is therefore essential for gene expression and cell viability. In contrast to the detailed mechanistic understanding of DNA repair pathways, cellular responses to RNA damage remain poorly defined^1,2^. This gap reflects a prevailing view of RNA as a transient molecule primarily destined for degradation rather than active repair.

Chemical modifications represent a common form of damage to both, DNA and RNA molecules. In particular, oxidation and alkylation of the nucleobases and the sugar-phosphate backbone cause chemically defined and characterized RNA lesions^2–4^. Other stresses, such as UV irradiation or aldehydes, cause RNA damage by inducing intra-or intermolecular cross-links^5^. These lesions can be detected and resolved by translation-coupled mechanisms when they occur in the mRNA^6–9^ or rRNA^10,11^. The resolution of these instances of chemical RNA damage does not necessarily involve direct RNA repair, but rather involve signaling by stalled ribosomes followed by degradation. Among the diverse forms of RNA damage, strand breaks pose a particular challenge because they generate truncated RNAs with chemically incompatible ends and disrupted structural integrity, impeding translation and other RNA-dependent processes. While some DNA break repair pathways hinge on the double-stranded architecture of DNA and or the presence of an intact reverse-complementary strand in template-based mechanisms^12^, such scenarios are not possible for breaks within RNA. Auto-hydrolysis, metal-dependent hydrolysis, RNases, oxidative stress and radiation are among known inducers of RNA backbone breaks^13–16^. Breaks within mRNAs lead to ribosome stalling at the truncated 3′ end, which triggers no-go decay (NGD)^17,18^. RNA strand breaks within rRNA have been linked to the ribotoxic stress response (RSR) that triggers translational arrest and apoptosis independently of DNA damage^19–21^. Both, NGD and RSR resolve RNA backbone breaks by degradation rather than repair of the damaged RNA.

A conserved class of enzymes capable of resolving such breaks is tRNA ligases, which catalyze the formation of a new phosphodiester bond between single-stranded RNA ends^22,23^. Divergent evolution yielded two distinct, non-homologous families of eukaryotic tRNA ligases, RTCB (in humans/metazoans) and Trl1 (in plants and fungi)^24,25^. Both tRNA ligase types catalyze the ligation of 5′-hydroxyl termini to 3′-phosphorylated ends, including 2′,3′-cyclic phosphates (2′,3′-cP) and 3′-monophosphates (3′-P). However, their biochemical mechanism and structural architecture are vastly different. RTCB-type ligases are GTP-dependent and catalyze direct ligation via activation of the 3′ end^23,26,27^. Human RTCB represents the catalytic core of the pentameric RNA ligase complex^23^. It further requires the protein activation factor Archease for multiple turn-over catalytic cycles^28,29^. In contrast, Trl1-type ligases are tripartite enzymes that harbor three distinct enzymatic activities in a single polypeptide chain^30^. In a multi-step reaction, the N-terminal ligase domain catalyzes the final ATP-dependent strand ligation via 5′ end activation^22,31^.

The canonical cellular roles of eukaryotic tRNA ligases include the ligation of exon halves during tRNA splicing and non-conventional mRNA splicing during the unfolded protein response (UPR)^22,23,32–35^. Both RNA processing mechanisms involve endonucleolytic removal of an intronic sequence followed by RNA structure-mediated exon-exon ligation^36^. Beyond their established roles in tRNA maturation and stress-induced mRNA processing, the human tRNA ligase RTCB has been implicated in specialized contexts such as transposon excision^37^ and programmable RNA ligation^38,39^. The growing range of tRNA ligase substrates suggests a broader RNA ligation capacity to restore damaged transcripts. In bacteria, the homologous RtcB ligase has been implicated in RNA break repair reactions of structured RNAs such as tRNA and rRNA^40–44^. However, it has remained unclear to what extent eukaryotic tRNA ligases function as general RNA repair enzymes that maintain transcriptome integrity.

Here, we investigate the mechanism and cellular role of tRNA ligase-mediated RNA break repair. Using complementary biochemical, cell biological and nanopore direct RNA sequencing approaches, we show that tRNA ligases promote repair of RNA strand breaks in human and yeast ribosomes, and confer tolerance to RNA damage-inducing stresses. Together, our findings reveal that eukaryotic tRNA ligases function as versatile safeguards of RNA integrity, supporting the maintenance of gene expression under conditions of RNA damage.

## RESULTS

### tRNA ligases repair toxin-cleaved SRL RNA breaks

We selected site-specific, endoribonuclease-induced cleavage of ribosomal RNA (rRNA) as a model system to study the RNA break repair capability of eukaryotic tRNA ligases towards ribonucleoprotein (RNP) complexes. To this end, we chose the α-sarcin-type ribotoxin restrictocin from *Aspergillus restrictus*, which specifically cleaves the highly conserved sarcin-ricin-loop (SRL) within 23-28S rRNA (Figures 1A and 1B)^45^. These rRNA cuts inactivate translational activity of ribosomes and are thus detrimental for the cell^46–48^. As is typical for α-sarcin-type endonucleolytic ribotoxins, restrictocin cleaves within the tetraloop of the SRL producing a ∼500-nt long 3′ fragment, the so-called α-fragment (Figure 1C)^47^. The resulting RNA backbone break harbors a 2′,3′-cP, which can be further hydrolyzed to a 3′-P, and a 5′-OH end at the cut site^48^, which represent bona fide tRNA ligase substrates. For all in vitro repair experiments, we used two biochemically distinct tRNA ligase systems: human RTCB with its activating factor Archease^28,29^ and fungal Trl1 (from *Chaetomium thermophilum*)^49–51^. First, we tested cleavage and repair using an isolated SRL oligoribonucleotide substrate and recombinantly expressed enzymes (Figure 1B). Incubation of the SRL substrate with restrictocin resulted in the expected cleavage fragments, which were re-ligated (i.e. repaired) by both tested RNA ligases, human RTCB and fungal Trl1 (Figures 1D and 1E). These results confirm toxin-cleaved SRL as a viable substrate for RTCB- and Trl1-type eukaryotic tRNA ligases.

**Figure 1:**
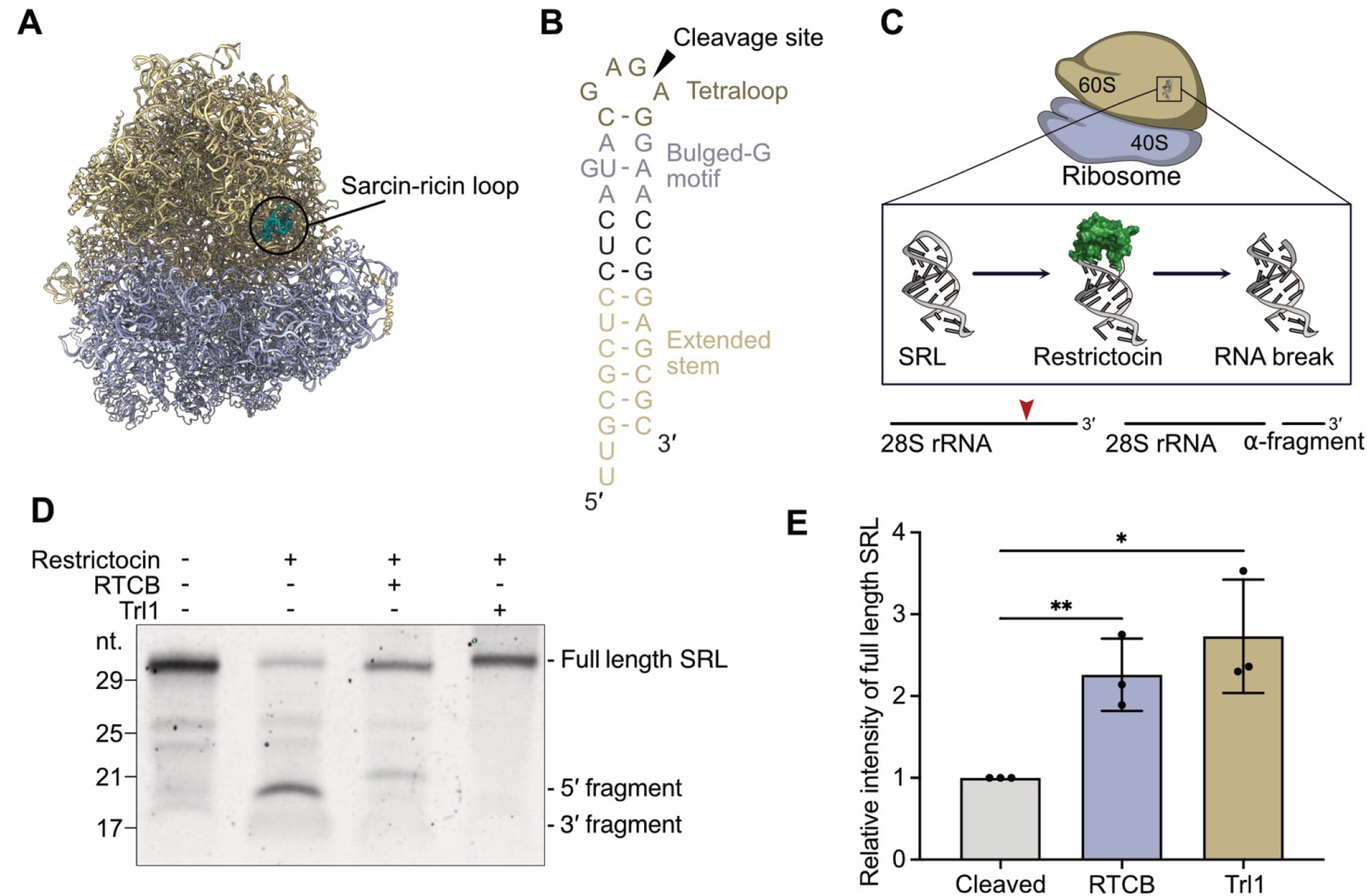
Repair of SRL-type oligonucleotide in vitro. (A) Structure of the human 80S ribosome with the SRL highlighted in teal. (B) Design of the SRL oligonucleotide with the universally conserved elements – the GAGA-tetraloop and the bulged-G motif – along with the cleavage site as well as the extended G-C rich stem are highlighted. (C) Schematic of 28S rRNA cleavage at the SRL by restrictocin and the formation of the 3′ α-fragment. (D) Urea-PAGE analysis of SRL oligonucleotide cleavage and repair. Incubation of SRL oligonucleotide with purified restrictocin leads to formation of the 3′ and 5′ fragment, which are reduced in intensity after incubation with either RTCB or Trl1 for 2 h. (E) Quantification of (D). Incubation of cleaved SRL-oligonucleotide with RTCB or Trl1 causes a significant increase in intensity of the full-length band intensity. n=3 independent experiments. Statistical significance was determined using two-tailed, unpaired t-test (defined significance levels: p<0.05 = *, p<0.01 = **, p<0.001 = ***, p<0.0001= ****). Error bars depict standard deviation (SD).

### Toxin-cleaved ribosomes are repaired by tRNA ligases

Next, we wanted to test if the same cleavage and repair mechanism also functions in the context of intact ribosomes. We hypothesized that efficient RNA repair of restrictocin-cleaved ribosomes would hinge on the α-fragment remaining bound to the ribosome, held in place by non-covalent interactions. To this end, we sedimented human ribosomes after cleavage by restrictocin and monitored the presence of the α-fragment by Northern blot analysis (Figure 2A). We found that the signal for the α-fragment was mostly present in the pellet fraction indicating continued association with ribosomes after cleavage (Figure 2B).

**Figure 2:**
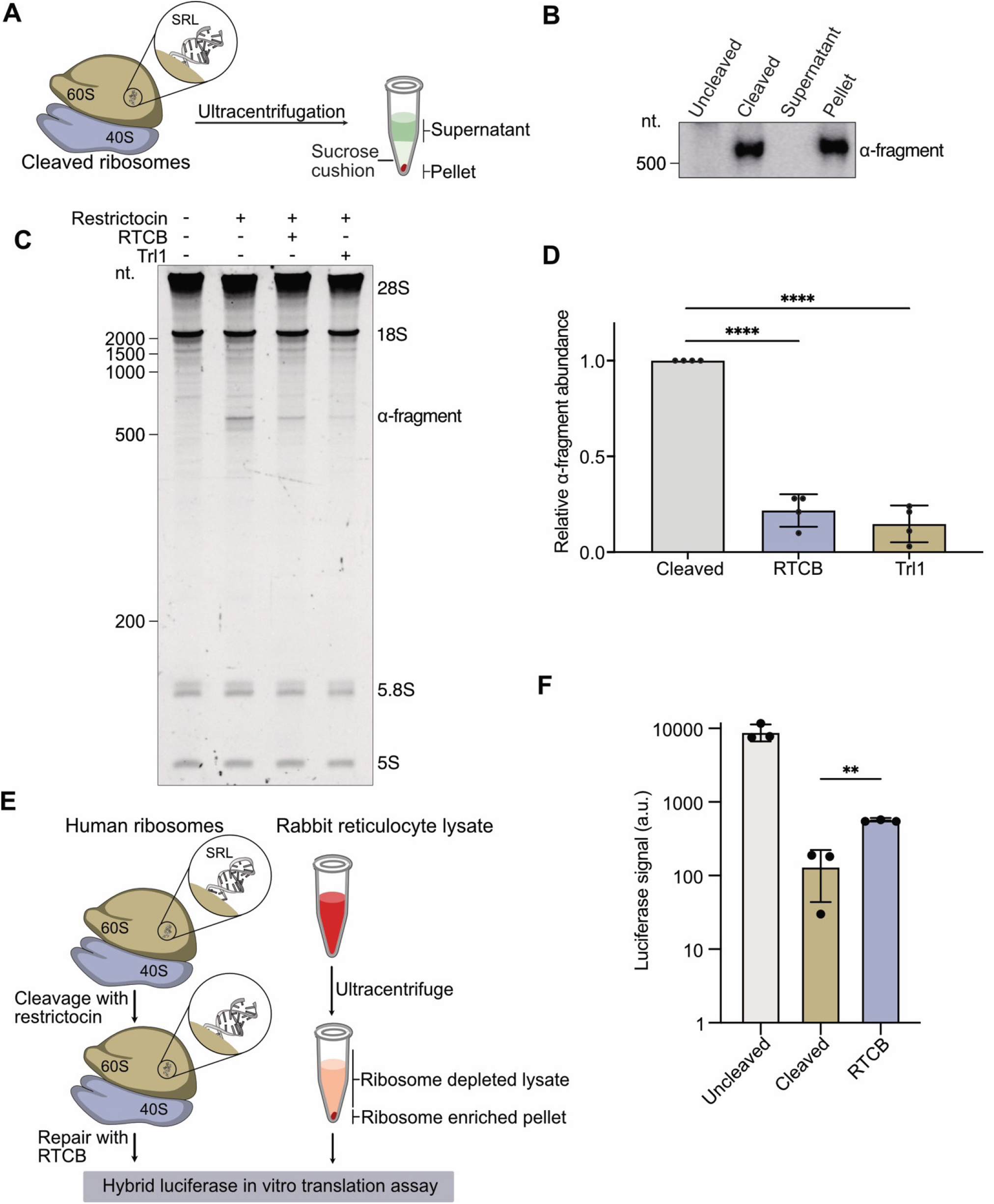
Repair of SRL in 80S ribosomes and translational reactivation in vitro. (A) Schematic of the density-ultracentrifugation assay to investigate whether the α-fragment remains attached to the ribosome after cleavage with restrictocin. (B) The α-fragment remains attached to the ribosome, by Northern blot it is only detected in the positive control and in the ribosome-containing pellet fraction. n=3 independent experiments. (C) Urea-PAGE analysis of 80S ribosome cleavage and repair. HEK293T 80S ribosomes are cleaved with purified restrictocin, after cleavage the α-fragment band appears, which is reduced in intensity after incubation with RTCB or Trl1 for 2 h. (D) Quantification of (C). Repair with both tRNA ligases leads to a significant decrease in α-fragment-band intensity, normalized to 5S rRNA as a loading control. n=4 independent experiments. (E) Schematic for the hybrid luciferase in vitro translation assay, using ribosome-depleted RRL and isolated 80S ribosomes from HEK293T cells to assay for translational activity after SRL cleavage and repair. (F) Luciferase activity measurement of hybrid luciferase in vitro translation assay. Repair of inactivated ribosomes with RTCB significantly increases translational activity in the hybrid luciferase in vitro translation assay. n=3 independent experiments. Statistical significances for panels in this figure were determined using two tailed, unpaired t-test (defined significance levels: p<0.05 = *, p<0.01 = **, p<0.001 = ***, p<0.0001= ****). Error bars depict standard deviation (SD).

Based on this observation, we tested if tRNA ligases are able to repair restrictocin-cleaved ribosomes. To this end, we monitored the amount of α-fragment using urea-PAGE analysis and Northern blot analysis of cleaved ribosomes in the presence of RTCB or Trl1. Incubation with either ligase resulted in a diminished signal for the α-fragment, indicating efficient repair of the cleaved SRL (Figures 2C, S1A and S1b). RTCB and Trl1 reduced the α-fragment intensity to 15% and 22%, respectively, indicating re-incorporation of the α-fragment and thus, rRNA repair in the majority of cleaved ribosomes (Figure 2D). Moreover, we showed rRNA repair of toxin-cleaved *S. cerevisiae* ribosomes using the fungal tRNA ligase Trl1 (Figures S1C, S1D, S1E and S1F). These findings establish ribosomes as a tractable model to study RNA break repair in structured RNPs.

### RNA break repair restores ribosome function

Because the SRL is essential for translation, we asked whether RNA break repair after insult restores ribosome function. The SRL is one of the most highly conserved sequences in 23-28S rRNA and is important for the interaction with elongation factors during translation elongation^52–54^. Cleavage of the SRL by α-sarcin-type nucleases causes translational arrest ^46,47^. In consequence, we postulated that ligase-mediated repair of restrictocin-cleaved ribosomes should result in functional restoration of ribosome activity and translational output. To this end, we established a hybrid in vitro translation (IVT) system based on rabbit reticulocyte lysates (RRL). We depleted endogenous ribosomes from the RRL by sedimentation, followed by addition of purified human ribosomes (Figure 2E). We first confirmed translational activity of the hybrid RRL IVT system using a luciferase reporter (Figure S1G). This luciferase-based hybrid IVT system allowed us to assess the impact of rRNA repair on active translation. We observed a significant increase in luciferase activity upon addition of RTCB-repaired human ribosomes compared to restrictocin-cleaved ribosomes, indicating concomitant repair of the SRL and translational re-activation (Figure 2F). Together, these results demonstrate that ligase-mediated repair can restore biological function of a damaged RNP following surgical cleavage of a single, critical phosphodiester bond.

### Direct sequencing quantifies RNA break repair at nucleotide resolution

To quantify the repair activity directed towards restrictocin-cleaved rRNA and to characterize the cleavage site with single nucleotide-resolution, we subjected untreated, toxin-cleaved and repaired ribosomes to nanopore direct RNA sequencing (DRS) (Figure 3A, scheme on the left). This long-read sequencing technology enabled simultaneous monitoring of the 28S rRNA breakage and repair at the single-molecule level. DRS enabled direct, nucleotide-resolution quantification of RNA break repair. RNA sequencing initiated from an artificially introduced polyA tail at the canonical 3′ end revealed a sharp drop in sequence coverage at the expected cleavage site, which was absent in the untreated samples (Figure 3A, scheme on the right, Figures S2A and S2B). Repair by either RTCB or Trl1 reverted this sequence coverage drop significantly (Figure 3B), resulting in an average repair efficiency of ∼60% for RTCB and ∼70% for Trl1, respectively (Figure 3C). We observed similar results for cleavage and repair of yeast ribosomes monitoring the coverage of 25S rRNA by nanopore sequencing (Figures S2C and S2D). Incubation with Trl1 yielded ∼60% repair of cleaved 25S rRNA (Figures S2E and S2F).

**Figure 3:**
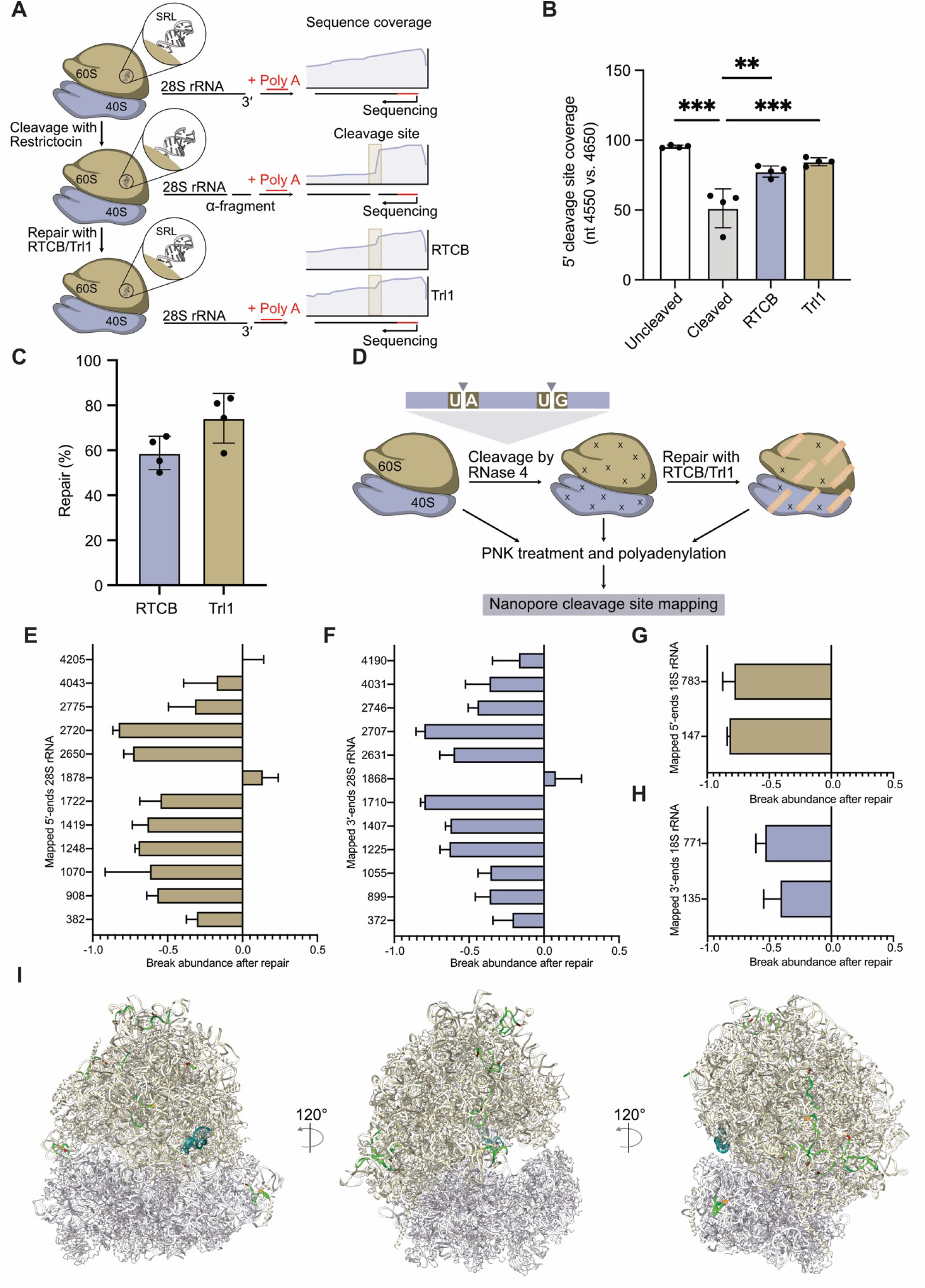
Direct RNA nanopore sequencing of RNA break repair. (A) Left: Schematic of the direct RNA nanopore sequencing for the restrictocin cleaved and RTCB/Trl1 repaired ribosomes. Right: Schematic of sequence coverage plots of the 28S rRNA around the SRL cleavage site. After cleavage with restrictocin, the sequence coverage drops, this drop is reduced after break repair with both tRNA ligases. (B) The sequence coverage drop is reduced by repair with both tRNA ligases. The sequence coverage 5′ of the cleavage site (nt 4550) was normalized to the sequence coverage 3′ of the cleavage site (nt 4650) for all analyzed samples after nanopore sequencing. n=4 independent experiments, statistical significance of cleavage and repair was determined by One-way ANOVA followed by Tukey’s multiple comparison test. (defined significance levels: p<0.05 = *, p<0.01 = **, p<0.001 = ***). (C) Around 60% of cleaved HEK293T ribosomes were repaired by RTCB, whereas ∼70% of ribosomes were repaired by Trl1. (D) Schematic of the unspecific breakage and repair assay, using RNase 4, which cleaves between any accessible U/A, or U/G dinucleotide on the isolated ribosomes, followed by repair with RTCB or Trl1 and nanopore sequencing-based breakage and repair site-mapping. (E-H) Most RNase4-induced RNA breaks show reduced relative 5′-end peak intensity or 3′-end peak intensity for both 28S and 18S rRNA after repair with RTCB (n=3 independent experiments). Hits were selected, when present in at least two replicates and when found by both analyses (<30 nt between 5′-end and 3′-end peak intensity). (I) Structural snapshots of the human ribosome (PDB 4ug0)^64^. Shown in dark violet is the SRL as a reference point, the breakage sites identified by either analysis are shown in dark red, the space in between the mapped break sites is shown in light green. Error bars depict standard deviation (SD).

### tRNA ligases repair diverse rRNA breaks dependent on local RNA context

Having established the repair of site-specific rRNA lesions, we next sought to determine whether tRNA ligases possess a broader repair capacity by introducing more widespread backbone breaks within rRNA. We selected RNase 4, which cleaves single-stranded RNA within uridine-purine dinucleotide sites (UA or UG), to introduce RNA breaks in intact ribosomes, followed by tRNA ligase-mediated repair (Figures 3D and S3A). Using DRS to map enriched 5′ ends, we identified multiple cleavage and repair sites across both the 18S and 28S rRNA (Figures S3B and S3C). Due to their short length, 5S and 5.8S rRNAs were excluded from this analysis. Comparison with identified 3′ ends allowed us to define multiple “high-confidence” repair sites with corresponding 5′ and 3′ ends (Figures 3E, 3F, 3G and 3H). When we mapped these sites onto the structure of the human ribosome, we found that the majority of cleavage events occurred within surface-exposed, unpaired segments of rRNA (Figure 3I and supplemental movie). We confirmed repair after RNase 4 cleavage in yeast ribosomes using fungal Trl1 ligase. Interestingly, the gel-based analysis of RNase 4-mediated break and repair of yeast ribosomes (Figure S3D) suggested more break-and-repair sites than those identified by nanopore sequencing (Figures S3E and S3F). Taken together, DRS enabled direct readout and quantification of rRNA repair by human and fungal tRNA ligase at both site-specific and non-specific endonucleolytic cleavage sites. Our data indicate surface exposure and structural accessibility on RNPs as key factors for efficient RNA ligase repair.

### RTCB levels correlate with RNA repair in cellulo

The results thus far showed the biochemical potential of eukaryotic tRNA ligases to repair rRNA breaks, but the experiments were performed in reconstitution systems. We next tested whether RTCB also repairs RNA strand breaks inside cells. We used siRNA-mediated knockdown (KD) to deplete endogenous RTCB in human HEK293T cells (Figure 4A). Introduction of purified restrictocin into these cells by lipofection-based delivery resulted in increased levels of α-fragment compared to WT (mock-treated) cells (Figure 4B). To assess whether RTCB ligase activity reflects the general RNA repair capacity in cells, we used a previously described split GFP trans-ligation reporter, employing self-cleaving ribozymes to produce mRNA fragments with compatible 2′,3′-cP and 5′-OH ends for ligase-mediated ‘stitching’ (Figure 4C). The inventors of this reporter suggested RTCB as the main catalytic driver behind mRNA trans-ligation^39^. We corroborated their findings by comparing split-GFP trans-ligation in WT versus RTCB KD cells. By fluorescence microscopy analysis, RTCB KD exhibited fewer GFP-positive cells compared to WT (Figure S4A). Flow cytometry analysis revealed a ∼2-fold reduction in GFP-positive cells upon RTCB KD (Figures 4D, S4B and S4C), and a significant decrease in mean GFP intensity among the GFP-positive cell population (Figure 4E and S4D) in the RTCB KD condition compared to WT. These results further substantiate RTCB as the main catalytic activity in trans-ligation of RNA fragments.

**Figure 4:**
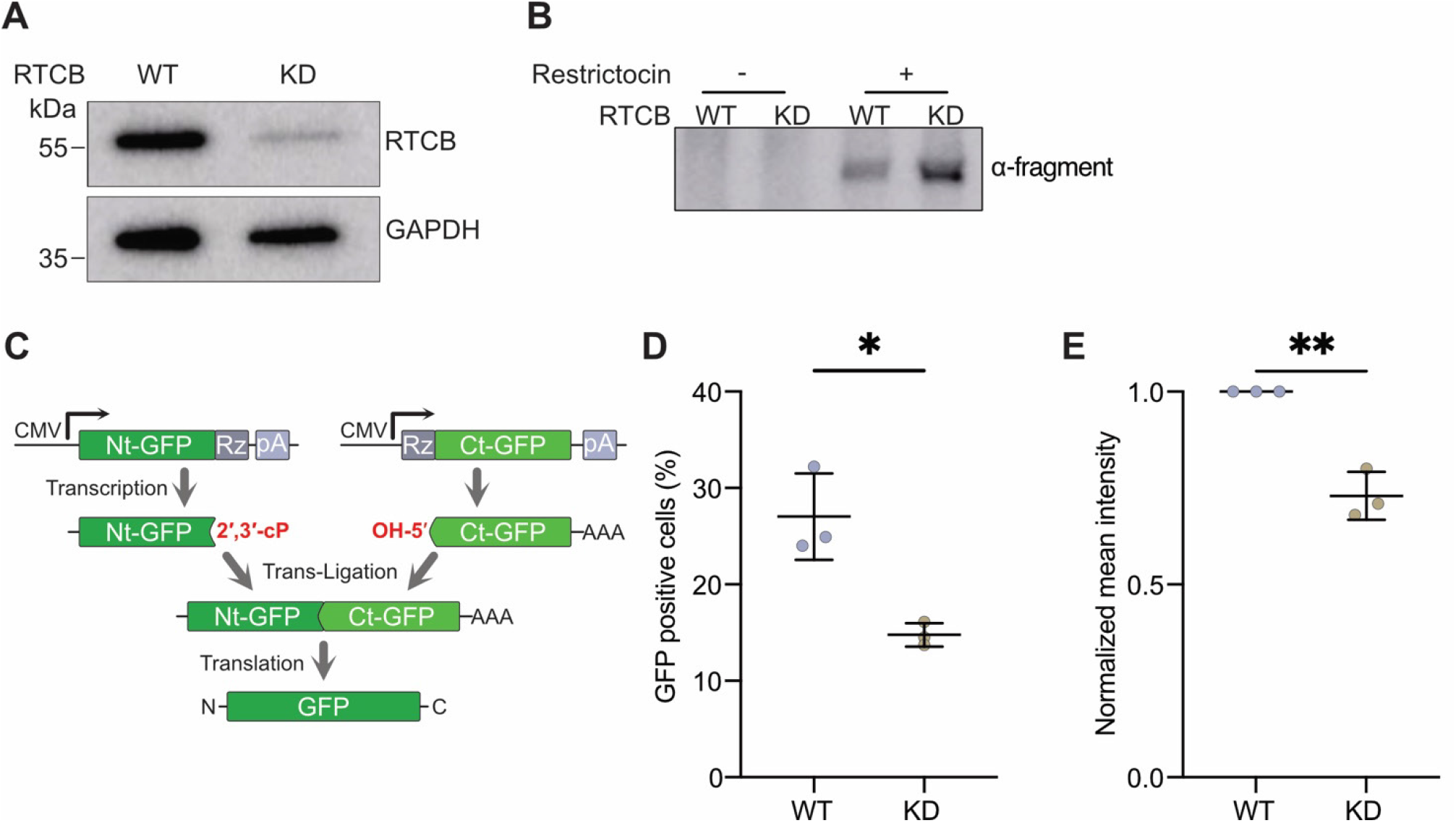
RTCB levels correlate with RNA repair in the cell. (A) Cellular RTCB protein levels are reduced after knockdown by siRNA, compared to cells, treated with mock-siRNA negative control (WT), as shown by western blot. (B) Northern blot of whole cellular RNA against the α-fragment-sequence. The α-fragment band intensity is stronger in RTCB KD compared to RTCB WT cells, 3 h after restrictocin was introduced to the cells via lipofection. n=3 independent experiments. (C) Schematic of the GFP trans-ligation co-transfection approach (adapted from Lindley et al. 2024 ^39^). (D) Flow cytometry analysis (488 nm) of RTCB WT and RTCB KD cells co-transfected with the trans-ligation reporters. Significantly less cells are GFP positive after co-transfection with GFP trans-ligation reporters in RTCB KD cells, compared to the WT cells. n=3 independent experiments. (E) Normalized GFP mean intensity is significantly reduced in GFP positive cells in the RTCB KD background vs. the RTCB WT cells. n=3 independent experiments. Statistical significances for panels in this figure were determined using two tailed, unpaired t-test (defined significance levels: p<0.05 = *, p<0.01 = **, p<0.001 = ***, p<0.0001= ****). Error bars depict standard deviation (SD).

### RTCB protects human cells from transcriptome-wide RNA damage

Finally, we explored the endogenous RNA break repair potential of cells. We devised an experimental setup that would allow us to trigger RNA backbone damage of the transcriptome combined with tunable repair capacity. A previous study reported induction of RNA breaks via the production of reactive oxygen species (ROS) by the vitamin K-analog menadione followed by capillary electrophoresis (Figure 5A)^55^. We adopted a similar strategy to monitor RTCB-mediated repair of RNA breaks in HEK293T cells. At increasing concentrations of menadione, RTCB KD cells showed stronger RNA fragmentation compared to WT cells (Figures 5B and 5C). RTCB overexpression (Figure S5A) provided further resistance to menadione-induced RNA breaks as monitored by capillary electrophoresis (Figure S5B). Increased rRNA fragmentation upon menadione treatment was also detectable by DRS. The cumulative read length for 28S and 18S rRNA shifted towards shorter reads upon menadione exposure, which was more pronounced in RTCB KD cells, indicating an excessive amount of RNA breaks in absence of RTCB (Figure 5D). As menadione-induced RNA breaks should not be limited to ribosomes, we determined the cumulative read length for other cellular RNPs using targeted nanopore sequencing. This analysis revealed a shift towards shorter reads in RTCB KD cells for *RN7SK*, a highly abundant structured small nuclear RNA^56,57^ and *RPPH1*, the RNA component of tRNA processing nuclease RNase P^58^ (Figure 5E). These findings extend RNA break repair beyond ribosomes and establish RTCB as a protector of multiple essential RNPs under stress conditions.

**Figure 5:**
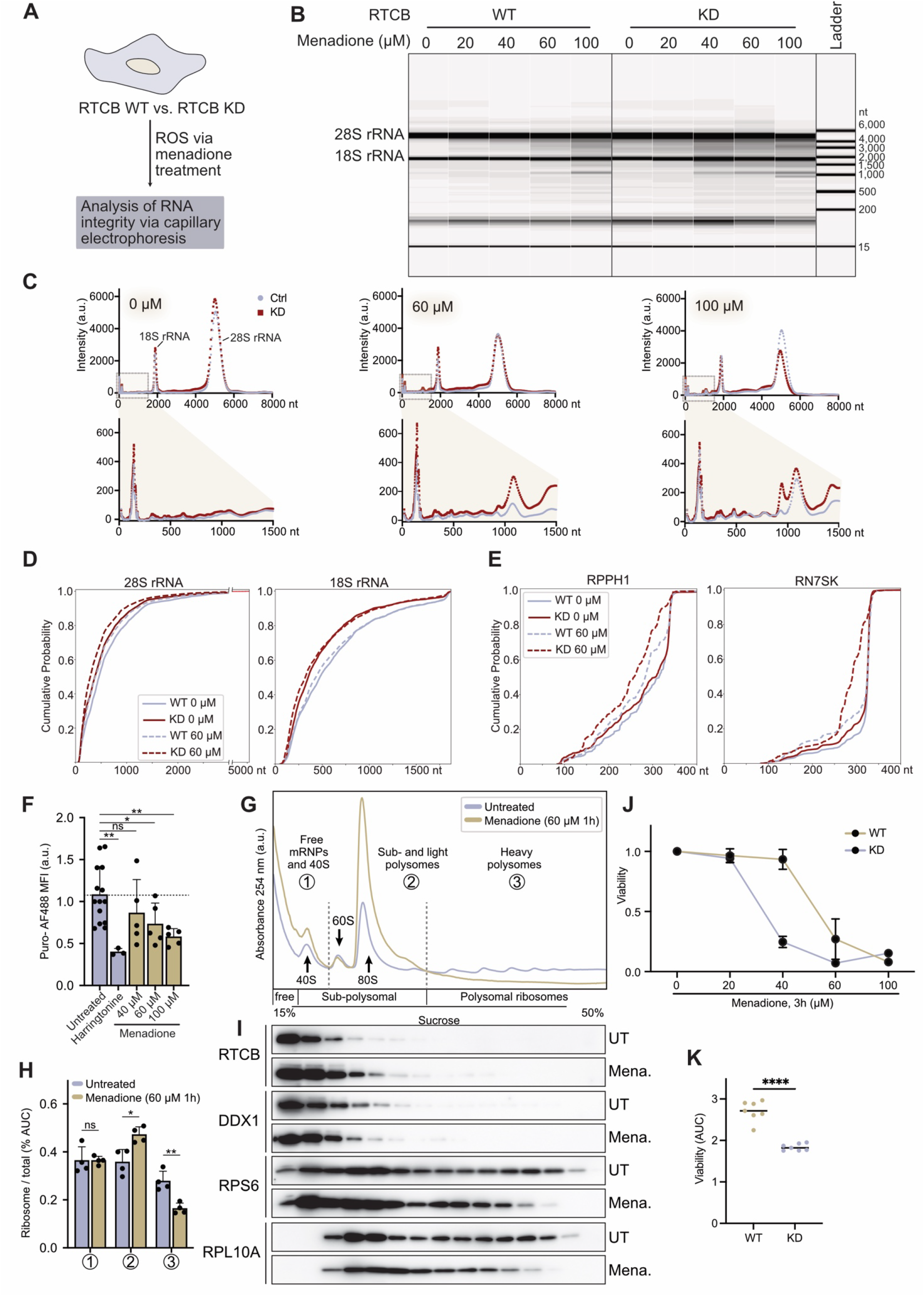
tRNA ligases protect cells from transcriptome-wide RNA damage. (A) Schematic of the experimental setup treating the RTCB knockdown cells with increasing concentrations of menadione for 3 h, before analyzing the isolated total cellular via capillary electrophoresis. (B) Electropherograms showing the effect of the RTCB knockdown with increasing concentrations of menadione. With increasing menadione concentrations, RNA is more fragmented in the RTCB KD compared to the WT. (C) Plots for the relative fluorescent units (a.u., arbitrary units) for three concentrations of menadione (0 µM, 60 µM and 100 µM) of the electropherograms shown in (B). Top: Whole cell RNA plot. Bottom: zoom in to small RNA fragments (nt 0-1500). (D) eCDF plots of cumulative probability for each read length after nanopore sequencing of RNA from cells treated with 0 µM and 60 µM menadione. Shown are read lengths for 28S rRNA (axis break between 2999 nt and 4500 nt) and 18S. (E) eCDF plots of cumulative probability for each read length after nanopore sequencing using specific sequencing adapters for *RPPH1* and *RN7SK* RNA from cells treated with 0 µM and 60 µM menadione. (F) Puromycin incorporation assay for HEK293T cells, treated with harringtonine as a control, and 3 different concentrations of menadione (40 µM, 60 µM, 100 µM) for 1 h. Median fluorescence intensity (MFI) of puromycin-AF488 is shown. For gating strategy refer to Figure S5C. (G) Polysome profiling (15-50% sucrose gradient fractionation) using either untreated cells, or cells treated with 60 µM menadione for 1 h. For quantification, profiles are categorized in three sections: 1) free mRNPs and 40S subunits, 2) sub- and light polysomes and 3) heavy polysomes. The peaks for 40S and 60S subunits, as well as for 80S ribosomes and polysomes are indicated. (H) Quantification of the AUC of the polysome profiles of ribosome/total ratios according to the 3 categories. N=4 independent experiments. (I) Western blot analysis specific for RTCB, and the ribosomal proteins RPLS6 (40S) and RPL10A (60S) using sucrose fractions from (H). (J) Viability assay of RTCB WT and RTCB knockdown cells treated with increasing concentrations of menadione. With increasing concentrations of menadione, cells are less viable in the RTCB KD compared to the WT. n=7 biological replicates. (K) Area under the curve (AUC) for the viability plots in (F). The AUC for the viability in the KD cells in significantly lower compared to the WT. Statistical significances for all panels in this figure were determined using unpaired t-test (defined significance levels: p<0.05 = *, p<0.01 = **, p<0.001 = ***, p<0.0001= ****). Error bars depict standard deviation (SD).

### RNA-damaging stress leads to translational shutdown and reduced cell viability

To study the cellular consequences of menadione treatment on cell viability in the context of RTCB levels, we first assessed the impact of increasing menadione concentrations on global translation in HEK293T cells. Due to the observed menadione-induced rRNA fragmentation (Figures 5B, 5C and 5D), we hypothesized that menadione causes a decrease in global translation. We used puromycin incorporation to measure nascent chain synthesis as a proxy for the cellular translation state. Compared to the translation initiation inhibitor harringtonine as a baseline for translation repression, we observed reduced global translation rates with increasing menadione concentrations compared to the untreated control (Figure 5F). To further analyze the translational status of menadione-treated cells, we performed sucrose gradient fractionation and polysome analysis in untreated and menadione-treated cells (Figure 5G). This analysis revealed a decrease in heavy polysomes (translating ribosome pool) and a respective strong increase in the sub- and light polysome fractions, particularly a large 80S ribosome peak, induced by treatment with 60 µM menadione for 1 h. This result indicates menadione-mediated global inhibition of translation (Figures 5G and 5H).

A previous interaction study identified the RNA helicase DDX1, a subunit of the human tRNA ligase complex, as a ribosome-associated protein (RAP)^59^. Here, we analyzed the association of DDX1 and RTCB with ribosomes based on their distribution along polysome gradients. We observed association of DDX1 but also RTCB with monosomal (80S) and subpolysomal ribosomes, which persisted upon menadione treatment (Figure 5I). These observations are consistent with a model in which RTCB transiently interacts with ribosomes and is thus positioned to repair their rRNA upon breakage to return them to the pool of translation-competent ribosomes.

Since we observed increased menadione-induced fragmentation for multiple essential RNA species upon reduced tRNA ligase levels in cells (Figures 5D and 5E), we next tested the impact of ligase-mediated repair on cell viability. When titrating menadione addition to WT and RTCB KD cells, we found a decrease in cell viability of the RTCB KD cells in presence of higher drug concentrations (Figures 5J and 5K). These findings suggest that tRNA ligase-mediated RNA repair contributes to stress tolerance maintaining overall translation and cell viability. Taken together, our data support a model in which RNA structure and RNP context enable repair of RNA strand breaks, providing an energy-efficient alternative to RNA decay.

## DISCUSSION

Cellular responses to RNA damage remain far less understood than DNA repair pathways. Exonucleolytic RNA decay pathways are widely considered the default when dealing with RNA strand breaks. Our findings challenge this view by demonstrating that eukaryotic tRNA ligases actively mend RNA strand breaks in vivo, establishing RNA break repair (RBR) as a conserved mechanism for maintaining RNA integrity. Both major classes of tRNA ligases, exemplified by human RTCB and fungal Trl1, promote repair of damaged RNAs in structured cellular contexts, including ribosomes and other essential ribonucleoprotein complexes.

Our data further suggest that RNA break repair is influenced by RNA architecture. Repair of toxin-induced and RNase-mediated breaks occurs within structured regions of rRNA and in the context of intact ribonucleoprotein assemblies, where RNA ends are likely retained in close spatial proximity. Consistent with this idea, we observe repair across multiple sites on the ribosomal surface and extend these findings to additional cellular RNPs, including the RNA component of the 7SK RNP and RNase P RNA. Together, these observations support a model in which RNA structure and RNP organization facilitate ligase-mediated rejoining of broken RNA ends. However, RTCB also repairs synthetic trans-ligation substrates, as in the presented split-GFP/ribozyme mRNA reporter, with unstructured ends^38,39,60^. Our findings expand the known catalytic repertoire of RTCB, demonstrating a vast repair spectrum that encompasses rRNAs, structured non-coding RNAs, and mRNAs across the human transcriptome.

The ability to repair RNA strand breaks has important implications for cellular stress responses. RNA damage, including RNA backbone breaks, has been linked to the ribotoxic stress response and can trigger translation inhibition and apoptosis independently of DNA damage ^19^. Our results show that tRNA ligase activity promotes tolerance to RNA-damaging conditions, such as oxidative stress, and enhances cellular viability. These findings suggest that RNA break repair mitigates the detrimental consequences of RNA damage and contributes to the maintenance of gene expression under stress conditions. In this context, RNA repair may act in competition with RNA decay pathways, which we and others proposed for fungal tRNA ligase Trl1^50,61^.

tRNA ligases are best known for their roles in tRNA splicing and unconventional mRNA processing during the unfolded protein response^23,32,33,62^. More specialized RNA repair reactions such as transposon-associated “SOS” repair and RNA ligation in engineered systems have been reported recently^37–39,60^. Our results suggest general RNA break repair as a broader and conserved function of eukaryotic tRNA ligases. In this expanded view, previously described ligase-dependent reactions represent specific instances of a more general capacity to restore RNA integrity following backbone cleavage.

Finally, our findings raise several important questions regarding the scope and regulation of RNA break repair. The extent to which different forms of RNA damage, including chemically induced lesions and endonucleolytic cleavage by endogenous RNases, are substrates for repair remains to be determined. In addition, recent identification of an alternative human RNA ligases, RLIG1, points to a potentially more complex network of RNA repair enzymes with distinct substrate preferences and cellular roles^55,63^. Elucidating how these activities are coordinated, and how RNA repair interfaces with stress signaling pathways such as the ribotoxic stress response, will be important areas for future investigation.

Together, our study establishes RNA break repair as a fundamental component of RNA quality control in eukaryotic cells and highlights tRNA ligases as central mediators of this process. By revealing that damaged RNAs can be restored, and not only degraded, our findings suggest that maintenance of RNA integrity relies on a dynamic interplay between repair and decay.

## METHODS

### Expression and purification of recombinant proteins

Recombinant human RTCB, human Archease and *Chaetomium thermophilum* Trl1 were expressed and purified as described previously^29,50^.

*Aspergillus restrictus* restrictocin (UniProt P67876) was expressed in *E. coli* BL21-CodonPlus (DE3)-RIPL (Agilent Technologies). Cells were grown in Luria broth (LB) medium containing kanamycin and 1% lactose at 30°C for 16 h and collected by centrifugation. The cells were lysed in lysis buffer (20 mM HEPES NaOH pH 7.5, 150 mM KCl, 1 mM MgCl_2_, 20 mM imidazole, 0.5 mM TCEP and 1x cOmplete protease inhibitor cocktail (Roche) using a microfluidizer. The first step of purification was performed by Ni^2+^ affinity chromatography using a 5 mL Ni-NTA column (Cytiva) in elution buffer (20 mM HEPES NaOH pH 7.5, 150 mM KCl, 1 mM MgCl_2_, 500 mM imidazole and 0.5 mM TCEP). The restrictocin containing fractions were pooled and concentrated using Amicon Ultra-15 Centrifugal Filter Units with a 10-kDa molecular weight cut-off (Merck), before being further purified through size-exclusion chromatography (SEC) on a HiLoad 16-600 Superdex 75 pg column (Cytiva). Restrictocin containing peak fractions were pooled, concentrated, flash-frozen in liquid nitrogen and stored at -80°C.

### Plasmid generation and cloning

All plasmids used in this study are listed in Table S1. For the generation of the pcDNA3.1(+)-3xFlag-*Hs*RTCB and pcDNA3.1(+)-3xFlag-*Ct*Trl1 plasmids, the *Hs*RTCB and *Ct*Trl1 sequences were cloned into an empty pcDNA3.1-3xFlag vector with standard restriction enzyme cloning, using the restriction enzymes EcoRI-HF and HindIII-HF (NEB) in CutSmart buffer (NEB) and primers listed in Table S2. The pET24A-ompA-Restrictocin-His6, pcDNA3.1(+)-NtGFP-Twister, pcDNA3.1(+)-HH-CtGFP were synthesized (BioCat).

### Polyacrylamide gel electrophoresis and Northern blotting

Samples in stop solution were unfolded at 80°C for 2 min and the reaction products were loaded onto a pre-run (300 V, 60 min) 6% or 15% urea gel. The gels were run in 1x TBE buffer at 150 V (constant voltage) at room temperature for 70 min. The gels were subsequently stained with SYBR Gold nucleic acid stain (Invitrogen, Life Technologies) to visualize RNA by ultraviolet trans-illumination. For northern blotting, RNA was transferred from Urea-PAGE gels to positively charged Nylon membranes (Roche) by semi-dry transfer at 100 mA for 45 min using the Trans-Blot SD (BioRad). Thereafter, RNA was crosslinked to the membrane with 1200 mJ/cm^2^ using the Stratalinker 1800 (Stratagene) pre-hybridized in hybridization buffer (5x SSC, 20 mM Sodium Phosphate buffer pH 7.2, 7% SDS, 2x Denhardt’s solution) at 55°C for 30 min and incubated with the Digoxigenin-labelled DNA probe (see Table S3) overnight in the hybridization oven. The membrane was washed twice in northern washing buffer 1 (2x SSC, 5% SDS) at 55°C, twice in northern washing buffer 2 (1x SSC, 1% SDS) at 55°C, once in northern washing buffer 3 (100 mM maleic acid, 150 mM NaCl, 0.3% Tween, pH 7.5) at RT and blocked in 10% block reagent (Roche) in maleic acid buffer (100 mM Maleic acid, 150 mM NaCl, pH 7.5). After blocking, the membrane was incubated with the anti-digoxigenin-AP fab fragments (Roche) for 30 min and subsequently washed thrice with northern washing buffer 3. Membranes were imaged with CDP-star chemiluminescence substrate (Roche) using the Amersham imager 600 (Cytiva).

### In vitro SRL oligonucleotide cleavage and repair

The SRL-type RNA oligonucleotide (Table S4) was diluted in restrictocin cleavage buffer (10 mM Tris HCl pH 7.4, 100 mM KCl, 1 mM MgCl_2_, 0.5 mM TCEP), thermally unfolded at 90°C for 1 min and refolded at room temperature by cooling down to ∼30°C. Cleavage was performed by adding 1 µM restrictocin-His for 60 min. After the cleavage, restrictocin-His was removed with nickel Sepharose™ beads (Cytiva) and membrane filtration (Corning®-Costar®-Spin-X® centrifuge tube filters). Ligation with RTCB was performed for 120 min at 30°C by adding 2 µM *Hs*RTCB, 4 µM *Hs*Archease, 10 mM GTP and 300 µM MnCl_2_ to the cleaved oligonucleotide. For the Trl1 ligation, 2 µM *Ct*Trl1, 10 mM ATP, 10 mM GTP and 3 mM MgCl_2_ was added for 120 min at 30°C. The reactions were quenched in 10-fold excess of stop solution (10 M urea, 0.1% SDS, 1 mM EDTA, trace amounts of xylene cyanol and bromophenol blue) and stored at -20°C. RNA samples were unfolded at 80°C for 2 min, analyzed by 15% denaturing urea-PAGE and visualized via SYBR™ Gold nucleic acid gel stain (Invitrogen, Life Technologies).

### Cell culture

Adherent HEK293T cells (CRL-3216, American Tissue Culture Collection, ATCC) (kind gift from the Christina Paulino lab) were grown at 37°C, 5% carbon dioxide and 70% humidity in 10cm dishes (Gibco) in Dulbecco’s modified Eagle’s medium (DMEM, Sigma-Aldrich), supplemented with 10% fetal bovine serum (FBS, Sigma-Aldrich) and antibiotic/antimycotic solution (Capricorn Scientific). In the Leppek lab, human HEK393T (ATCC: CRL-3216) were cultured in Dulbecco’s Modified Eagle’s Medium (DMEM, Gibco) containing 2 mM L-glutamine, supplemented with 10% fetal calf serum (Gibco), 100 U/ml penicillin and 0.1 mg/mL streptomycin (EMD Millipore, or Gibco,) (regular medium, RM) at 37°C in 5% CO_2_-buffered incubators.

### Human ribosome isolation

The human ribosome purification protocol was adapted from the Klaholz lab^65^. HEK293T cells were lysed in lysis buffer (50 mM HEPES KOH pH 7.5, 0.5% NP40, 6 mM MgOAc, 300 mM KOAc, 1 mM TCEP, 1x cOmplete protease inhibitor cocktail (Roche), RNasin® Plus RNase inhibitor [Promega]). Lysates were cleared from cell debris by centrifugation, the supernatant was loaded on a 30% sucrose cushion (20 mM HEPES KOH pH 7.5, 2 mM MgOAc, 500 mM KOAc, 30% sucrose, 1 mM TCEP, 1x cOmplete protease inhibitor cocktail (Roche), RNasin® Plus RNase inhibitor [Promega]) and centrifuged at 116,000 g for 16 h at 4°C (70Ti rotor, 39,000 rpm). The pellet was resuspended in resuspension buffer (20 mM HEPES KOH pH 7.5, 6 mM MgOAc, 150 mM KOAc, 6.8% sucrose, 1 mM TCEP, RNasin® Plus RNase inhibitor [Promega]) and subsequently treated with 1 mM puromycin and 1 mM GTP at 37°C. After a 1 min clearing spin at 13,000 g, the supernatant was layered over a 15%-50% sucrose gradient in gradient buffer (20 mM HEPES KOH pH 7.5, 2 mM MgOAc, 150 mM KOAc, 1 mM TCEP, 1x cOmplete protease inhibitor cocktail (Roche), RNasin® Plus RNase inhibitor [Promega]) and centrifuged at 60,000 g for 16 h at 4°C (SW28 rotor, 21,00 rpm). Fractions were taken from the top and the absorption at 260 nM was measured using a NanoPhotometer® (Implen). Peak fractions containing 80S ribosomes were pooled and loaded on a 30% sucrose cushion and centrifuged as described previously. The pellet was resuspended in ribosome storage buffer (20 mM HEPES KOH pH 7.5, 5 mM MgOAc, 100 mM KOAc, 1 mM TCEP), flash-frozen in liquid nitrogen and stored at -80°C.

### Yeast growth conditions and yeast ribosome isolation

*S. cerevisiae* W303 cells were grown in YPD medium at 30°C, 120 rpm to an OD_600_ of 1. Cells were harvested by centrifugation, resuspended in storage buffer (20 mM HEPES KOH pH7.5, 5 mM MgOAc, 50 mM KOAc, 2 mM DTT, 1x cOmplete protease inhibitor cocktail [Roche]) and frozen dropwise in liquid nitrogen. Lysis was performed by cryo-milling (Retsch), lysate was cleared by centrifugation. The supernatant was loaded on a 30% sucrose cushion (20 mM HEPES KOH pH 7.5, 5 mM MgOAc, 500 mM KOAc, 2 mM DTT, 1x cOmplete protease inhibitor cocktail [Roche]) and centrifuged at ∼200,000 g for 16 h at 4°C (70 Ti rotor, 55,000 rpm). The pellet was resuspended in ribosome storage buffer and treated with 1 mM puromycin and 1 mM GTP at 37°C. After a 1 min clearing spin at 13,000 g, the supernatant was layered over a 15%-40% sucrose gradient in gradient buffer (20 mM HEPES KOH pH 7.5, 5 mM MgOAc, 150 mM KOAc, 2 mM DTT, 1x cOmplete protease inhibitor cocktail [Roche]) and centrifuged at ∼60,000 g at 4°C (SW28 rotor, 21,000 rpm). The gradient was fractionated and the absorption at 260 nM was measured using a NanoPhotometer® (Implen). Peak fractions were pooled, and ribosomes were precipitated with 7% w/v PEG 20,000. Ribosomes were flash-frozen in liquid nitrogen and stored at -80°C.

### α-fragment attachment assay

500 nM of ribosomes isolated from HEK293T cells were incubated with 100 nM of purified restrictocin-His in ribosome storage buffer at 37°C for 30 min, before being layered over a 30% sucrose cushion (20 mM HEPES KOH pH 7.5, 2 mM MgOAc, 500 mM KOAc, 30% sucrose, 1 mM TCEP, RNasin® Plus RNase inhibitor [Promega]) and centrifuged for 2 h at ∼200,000 g at 4°C (TLA100 rotor 45,900 rpm). Thereafter, the supernatant and pellet fractions were resuspended in 1 mL of Trizol (Invitrogen, Life Technologies) and the ribosomal RNA was purified using the RNA clean and concentrator-25 kit (Zymo Research). RNA concentrations were measured using a NanoPhotometer® (Implen). 1/30 of the isolated RNA from each condition was diluted in 30-fold excess of stop solution (10 M urea, 0.1% SDS, 1 mM EDTA, trace amounts of xylene cyanol and bromophenol blue) and analyzed by northern blotting.

### In vitro ribosome cleavage and repair with RTCB or Trl1

To cleave both yeast and human ribosomes with restrictocin, 500 nM of ribosomes were incubated with 100 nM of restrictocin-His in ribosome storage buffer at 37°C for 30 min, before restrictocin-His was removed with nickel Sepharose™ beads (Cytiva) and membrane filtration (Corning®-Costar®-Spin-X® centrifuge tube filters).

For the cleavage with RNase 4, 500 nM of both *s*.*c*. and *h*.*s*. ribosomes were treated with 1% (v/v) RNase 4 (NEB) in NEBuffer™ 1.1 (NEB). After 10 min, the reaction is stopped by addition of murine RNase inhibitor (NEB) for 10 min.

Ligation with RTCB was performed for 120 min by adding 2 µM *Hs*RTCB, 4 µM *Hs*Archease, 10 mM GTP and 300 µM MnCl_2_ to the cleaved ribosomes. For the Trl1 ligation, 2 µM *Ct*Trl1, 10 mM ATP, 10 mM GTP and 3 mM MgCl_2_ was added for 120 min. The reactions were stopped by adding 1 mL of Trizol (Invitrogen, Life Technologies) and the ribosomal RNA was purified using the RNA clean and concentrator-25 kit (Zymo Research). RNA concentrations were measured and normalized using a NanoPhotometer® (Implen). RNA was either diluted in 30-fold excess of stop solution for analysis via polyacrylamide gel electrophoresis and Northern blotting, or flash frozen in liquid nitrogen and stored at -80°C for further analysis via nanopore sequencing.

### Hybrid luciferase in vitro translation assay

Hybrid in vitro translation protocol was adapted from the Iwasaki lab protocol^66^. Rabbit reticulocyte lysate (Promega) was centrifuged at ∼200,000 g for 2 h at 4°C (TLA100 rotor 45,900 rpm) and the supernatant was taken off, aliquoted, flash-frozen in liquid nitrogen and stored at -80°C. For the reactions, human ribosomes were cleaved with restrictocin and repaired by RTCB as described previously. Thereafter, ribosomes were purified from the repair reaction by dilution in ribosome storage buffer and concentrated with 100 kDa MWCO Amicon centrifuge filters (Millipore). In vitro translation reactions were performed with ribosome-depleted RRL, 50 nM either uncleaved, cleaved, or repaired HEK293T ribosomes, and in vitro translation master mix (75 mM KCl, 0.75 mM MgCl_2_, 10 µM amino acid mixture minus Leucin (Promega), 10 µM amino acid mixture minus Cysteine (Promega), 1 µg Luciferase control RNA [Promega]) at 30°C for 120 min. Luciferase signal was measured using the luciferase assay system (Promega) at the GloMax® 20/20 luminometer (Promega). Blank measurement was performed with the reaction without HEK293T ribosomes and subtracted from the subsequent samples.

### Pre-treatment of RNA for nanopore direct RNA sequencing

RNA from *in vitro* cleavage assays were subjected to T4 polynucleotide kinase (PNK) treatment to generate ends compatible with polyA tailing and/or direct RNA-sequencing (5’ phosphate and 3’-OH).

1000 ng RNA was incubated in 1x T4 PNK buffer, 1 mM ATP and 10 U T4 PNK (New England Biolabs) for 30 min at 37°C. The enzyme was denatured by a 20 min incubation at 60°C. Reactions were cleaned up using the RNA Clean & Concentrator-5 kit (Zymo Research).

RNA that was T4 PNK treated (RNase 4 assays, 5’ fragment analysis of restrictocin cleavage, total RNA from menadione treated cells) or not (restrictocin assays) were polyadenylated using a poly(A) tailing kit (Thermo Scientific). Reactions containing 500 ng RNA, 1x E-PAP buffer, 1 mM ATP, 5 mM MnCl_2_, 40 U RiboLock (Thermo Scientific) and 2 U E-PAP were incubated 5 min at 37°C. Reactions were cleaned up using the RNA Clean & Concentrator-5 kit (Zymo Research).

### Nanopore direct RNA sequencing

Direct RNA sequencing (DRS) was carried out using a custom protocol with WarpDemuX barcode adapters ^67^ and the SQK-RNA004 kit (Oxford Nanopore Technologies). For targeted sequencing of *RPPH1* and *RN7SK*, WarpDemuX-rRNA (https://github.com/KleistLab/WarpDemuX, Naarmann-de Vries *et al*., under review [“WarpDemuX-rRNA -Multiplexed sequence-specific direct RNA sequencing of ribosomal RNA and beyond”]) was used for demultiplexing. The RPPH1 adapter is described in Naarmann-de Vries et al., under review and the *RN7SK* adapter sequence has been described by Leger et al. 2021^68^. Custom barcode adapters (either targeting the polyA tail or the 3′ end of *RPPH1* and *RN7SK*) were annealed as described previously^69^. 200 ng RNA in a total volume of 10.5 µl were combined with 3 µl Quick ligation buffer (New England Biolabs), 0.5 µl barcoded adapter and 1 µl T4 DNA ligase (high concentration, New England Biolabs). Ligation was carried out for 10 min at room temperature. A reverse transcription mixture composed of 14 µl nuclease-free water, 8 µl 5x Induro buffer and 2 µl 10 mM dNTPs (Thermo Scientific) was added and mixed by pipetting before addition of 1 µl Induro RT. Reverse transcription reactions (to dissolve secondary structures) were carried out for 30 min at 60°C, followed by enzyme denaturation (10 min, 70°C). 2 µl 0.5 M EDTA were added prior to pooling of all barcoded samples (3-plex to 4-plex) and reactions were cleaned up with 1.8x RNA Clean XP beads (Beckman Coulter). After two wash steps with 80% ethanol, elution was performed in 23 µl nuclease-free water. The RLA sequencing adapter was subsequently ligated as described in the manufacturer’s protocol. Final libraries were eluted in 33 µl REB. The concentration of the final libraries was determined on a Qubit fluorometer using the dsDNA HS kit (Thermo Scientific). Libraries were loaded completely on PromethION RP4 flow cells (Oxford Nanopore Technologies) and sequenced on a PromethION P24-A100 device (Oxford Nanopore Technologies) equipped with MinKNOW v25.03.7 to v25.09.16 with the following setting: 500 Mb estimated bases as data target, high accuracy basecalling (Dorado basecall server v7.8.3 to v7.11.2 (using v5.1 RNA004 models), Q score threshold of 7.)

Reads were demultiplexed using the WarpDemuX workflow^67^ and aligned to a human (NR_003286.4, NR_003287.4) or yeast rRNA reference with minimap2. Data were visualized with Integrative Genomics Viewer (IGV).

### Nanopore data analysis of restrictocin *in vitro* cleavage

The restrictocin-induced cleavage and RTCB-or Trl1-mediated repair was analyzed by different means. For the “coverage drop” analysis, the coverage upstream and downstream of the cleavage site was derived from IGV. For human samples, positions 4550 and 4650 were assessed. For yeast samples, positions 3000 and 3100 were analyzed. Statistical significance of cleavage and repair was determined by One-way ANOVA followed by Tukey’s multiple comparison test (human: n=4, yeast: n=3, defined significance levels: p<0.05 = *, p<0.01 = **, p<0.001 = ***). Furthermore, the relative fraction of 5’ read ends per position was derived from the mapped bam files using a custom python script. 5’ read end peaks with a height of 0.1 in a 20 nt distance were considered, if present in the respective cleaved samples and consistently detected among replicates. Cleavage and repair at the identified cleavage site (3040 for yeast, 4628 for human) was analyzed as described above.

### Nanopore data analysis of RNase 4 *in vitro* cleavage

3’ read ends and 5’ read ends were quantified as described above. Consistently detected peaks (present in at least 2 replicates) in cleaved samples were considered for further analysis. Read lengths of the individual rRNAs were plotted as empirical cumulative distribution function (eCDF) in Python.

### Transient transfections and siRNA mediated knockdowns

HEK293T cells were transfected using Lipofectamine™ 3000 (Thermo Scientific) according to the manufacturer’s protocol. 16 h after transfection, cells were used for one of the following: protein lysate preparation, using M-PER™ Mammalian protein extraction reagent (Thermo Scientific), RNA extraction, using Trizol (Invitrogen, Life Technologies) and RNA clean- and concentrator kit (Zymo Research), or fixation, followed by widefield microscopy. For co-transfections, plasmids were used in equimolar concentrations. SiRNA mediated knockdowns were performed with Silencer® siRNAs (siRNA ID:124763, Thermo Scientific) and Lipofectamine™ RNAiMAX (Thermo Scientific) according to the manufacturer’s protocol for 72 h. Negative control (RTCB WT) cells were treated in the same way as the KD cells, and mock-transfected with the same amount of Ambion™ Silencer™ Negative Control Nr. 1 siRNA (Invitrogen™).

### Widefield microscopy and flow cytometry

For widefield microscopy, cells were reverse transfected with the split-GFP constructs in Collagen IV coated µ-Slide 8-well plates (Ibidi). After 24 h, cells were washed with 1x PBS (Gibco), fixed in 4% paraformaldehyde (Thermo Scientific) in 1x PBS, then washed three times with 1x PBS and imaged by widefield microscopy using 20x magnification (Eclipse Ti2 series, Nikon).

For flow cytometry, cells were co-transfected with the split-GFP constructs in a 6-well plate format. After 24 h, cells were washed with 1x PBS, fixed in 4% paraformaldehyde (Thermo Scientific) in 1x PBS, then washed three times in 1xPBS and analyzed with the BD FACSymphony A1 (Excitation laser: 488 nm, Detection Filters GFP 505LP and 530/30, Software BD FACSDiva 9, Threshold FSC-H set to 5000) The cell population was gated based on FSC-A and SSC-A. Single cells were gated based on FSC-A vs. FSC-H (fig. S4B). Both percentage of GFP-positive cells and the mean GFP signal for the GFP-positive cells was determined.

### Restrictocin lipofection

HEK293T WT and RTCB KD cells were prepared as described previously. For the lipofection of one well in a 24-well plate (Greiner), Pierce™ Protein transfection reagent (Thermo Scientific) was resuspended in 0.25 µg of restrictocin, diluted in 1x PBS. Cells were washed once with 1x PBS and then the protein transfection mix was added to the cells in 250 µL serum-free Opti-MEM (Gibco). Cells were incubated for 3 h before cellular RNA was extracted using Trizol (Invitrogen, Life Technologies) and RNA clean and concentrator-25 (Zymo Research), according to the manufacturer’s protocols. RNA concentrations were measured at the Nanophotometer (Implen). The same amount of RNA for each sample was diluted in 30-fold excess of stop-solution and further analyzed by northern blotting.

### Menadione treatment and analysis of RNA integrity

HEK293T cells with a siRNA mediated RTCB knockdown were treated with increasing concentrations of menadione (0 µM, 20 µM, 40 µM, 60 µM, 100 µM), from 1,000x stocks diluted in ethanol^55^. After 3 h of incubation, total RNA was isolated using Trizol (Invitrogen, Life Technologies) and RNA clean and concentrator-25 (Zymo Research), according to the manufacturer’s protocols. Total RNA from menadione treated cells was analyzed on a Fragment Analyzer device (Agilent Technologies) employing the standard RNA kit.

Total RNA from control and RTCB-depleted cells treated with 60 µM menadione or untreated, was subjected to T4 PNK treatment, poly(A) tailing and Nanopore DRS of rRNAs. Alternatively, *RPPH1* and *RN7SK* were targeted with sequence-specific adapters for DRS (see above). Median read lengths of the individual rRNAs were plotted as eCDF in Python.

Cellular viability was measured using the CellTiterGlo 2.0 assay system (Promega) according to the manufacturer’s protocol in 96-well format.

### FACS-based puromycin incorporation assay

For fluorescence-based FACS readout of puromycin incorporation, we adapt the SCENITH (single-cell energetic metabolism by profiling translation inhibition) protocol^70^. For this, 75,000 HEK393T cells were seeded per well in 96-well U-bottom plates and cells were treated the following day with 0 µM, 40 µM, 60 µM, or 100 µM menadione (Roth, in EtOH) for 1 or 3 h at 37°C before fixation. Time points were timed so that cells were harvested together. Experiments were performed in triplicates with technical repeats in triplicates. Untreated cells and cells pre-treated with harringtonine (2 mg/mL 1,000x stock, final 2 μg/mL; Abcam) (translation initiation inhibitor) for 15 min at 37°C were included as negative controls. Cells were then incubated with puromycin dihydrochloride (*Streptomyces alboniger*, Sigma). Final 2x puromycin (20 mg/mL 2,000x stock, final 10 μg/mL) was added to each well for a final volume of 120 μL per well. Each well was mixed and incubated for 30 min at 37°C in the incubator. The plate was transferred on ice and ice-cold FACS buffer (1x PBS, 2% FCS, 2 μM EDTA) was used to wash the cells. 100 µL per well of FACS buffer was added, the plate was centrifuged and buffer was removed by inverting the plate forcefully once or by pipetting the buffer off. Plates were centrifuged at 1,500 rpm for 4 min at 4°C and supernatant was discarded. Cells were resuspended in 50 μL Live/Dead reagent (Zombie, UV) at 1:400 dilution in 1XPBS (Thermo Fisher Scientific) and FC block at 1:200 dilution in 1x PBS and the plate was incubated for 15 min at 4°C in the dark in the refrigerator. The plate was centrifuged at 1,500 rpm for 4 min at 4°C and the supernatant was discarded. The surface staining panel (peritoneal macrophages: CD19 (#29), F4/80 (#201), C11b (#137) at 1:200 dilution in 1XPBS was added and the plate was incubated for 25 min at 4°C in the dark in the refrigerator. The cells are washed with ice-cold FACS buffer, centrifuged at 1,500 rpm for 4 min at 4°C and the supernatant was discarded. Cells were resuspended in 100 μL Foxp3 Fixation/Permeabilization solution (Foxp3 permeabilization wash kit (Thermo Fisher; brown concentrate: 1/4, White dilute ¾)) and the plate was incubated for 30 min at RT protected from light. The plate was washed twice with 100 μL per well of 1x Permeabilization Wash Buffer (10x stock is diluted to 1x in Ampuwa water). 30 μL per well of puromycin-antibody solution (1:1,000 dilution of the stock in 1x Permeabilization Wash Buffer) is added and the plate was incubated for 25 min at 4°C protected from light in the refrigerator. 70 μL of 1x Permeabilization Wash buffer was added for a final volume of 100 µL, the plate was centrifuged at 1,500 rpm for 4 min at 4°C, and the supernatant was discarded. For FACS measurement, pellets of stained cells were resuspended in a volume of 100 μL FACS buffer. The puromycin signal intensity was measured by using the FACS cytometer ID7000 5L Spectral Cell Analyzer (Sony). FACS data were exported as FCS files and further analyzed by FlowJo v10.8 (BD). Viable cells and singlets were identified based on the FSC and SSC channels. The puro signal was measured and used for calculation of the median fluorescence intensities (MFIs).

### Sucrose gradient fractionation analysis

For sucrose gradient fractionation of HEK293T cell lysates, ∼1.0-1.2 × 10^6^ HEK393T cells were seeded into 15-cm dishes in RM per sample. The next day, cells at 50-60% density were treated with 0 and 60 µM menadione (Roth, 1T96.1, in EtOH) in RM for 1 hour at 37°C prior to harvest. For cycloheximide (CHX) (Sigma-Aldrich) treatment, HEK293T cells were incubated in RM with 100 µg/mL CHX for 3 min at 37°C in the incubator, washed once with cold DPBS (Thermo Fisher Scientific) with CHX, and harvested with 0.05% Trypsin-EDTA (Thermo Fisher Scientific) with CHX at 37°C. The reaction was stopped by adding RM+CHX. After centrifugation of cells at 1,000 g for 3 min at RT, cells were washed with 1 mL DPBS+CHX, cells were transferred into 1.5 mL DNA LoBind safe-lock tubes (Eppendorf) and incubated a total of 20 min on ice from the CHX treatment start. Cells were collected by centrifugation at 1,000 g at 4°C. The DPBS was removed and the cell pellet was flash frozen in liquid nitrogen and stored at -80 for later polysome analysis. For polysome analysis, cell pellets were lysed in polysome lysis buffer (25 mM Tris-HCl pH 7.5 (Ambion,), 150 mM NaCl (Ambion), 15 mM MgCl_2_ (Ambion), 1 mM DTT (Ambion), 8% glycerol (Sigma-Aldrich), 1% Triton X-100 (Sigma-Aldrich), 0.1 mg/mL cycloheximide (CHX) (Sigma-Aldrich), 100 U/mL SUPERaseIn RNase Inhibitor (Ambion), 25 U/mL TurboDNase (Ambion), Mini complete Protease Inhibitor EDTA-free (Sigma-Aldrich), 1% sodium deoxycholate, in nuclease-free water [Thermo Fisher Scientific]). For ∼1.0-1.2 × 10^6^ HEK393T cells, 400 µL of lysis buffer was used to lyse the cells by vortexing each sample for 30 s followed by 30 s incubation on ice, and the lysate was forced 6x through a syringe with a 30G needle. Lysis continued for 30 min on a rotator at 4°C and vortexing 30 seconds 3x every 10 min. After lysis, nuclei were removed by centrifugation at 10,000 g for 10 min at 4°C. Supernatants were transferred to a new 1.5 mL DNA LoBind tube (Eppendorf). RNA concentrations were measured using a Nanodrop UV spectrophotometer (Thermo Fisher Scientific) for normalization of RNA amounts in the lysates. The clarified cytoplasmic extracts were layered onto a linear 15-50% sucrose gradient (15%-50% sucrose (Calbiochem, OmniPur Sucrose) (w/v), 25 mM Tris-HCl, pH 7.5, 150 mM NaCl, 15 mM MgCl_2_, 1 mM DTT, 100 mg/mL CHX in nuclease-free water) in open-top polyclear centrifuge tubes (Seton) and centrifuged in a SW41Ti rotor (Beckman) for 2 h at 40,000 rpm at 4°C in a Beckman ultracentrifuge (Optima XPN-100). Fractions were collected by the Gradient Station ip System (BioComp Instruments) with continuous A_260_ measurement. After collection of polysome fractions in 2 mL safe-lock tubes (Eppendorf), samples were frozen on dry ice and stored at -80°C. We quantified the area under the curve between free fraction, light and heavy polysomes to assess ribosome association of a protein based on its distribution in the gradient.

### Bead-based protein extraction from polysome fractionation samples

For bead-based purification of total protein from sucrose fractions, fractions were thawed on ice. For isolation of total protein from fraction samples the StrataClean Resin (Agilent Technologies, 400714) was used. For washing the beads, 10 µL of 50% bead slurry per sample (15 samples per gradient) were washed with 1x Polysome Buffer (without sucrose) (25 mM Tris pH 7.5 (Ambion), 150 mM NaCl Ambion), 15 mM MgCl_2_ (Ambion), 1 mM DTT (Ambion), in nuclease-free water [Thermo Fisher Scientific]) in a 2 mL protein LoBind tube (Eppendorf). Beads were washes twice with 1 mL 1x Polysome Buffer and centrifuged for 1 min at 3,000 rcf at 4°C. After the second wash step, supernatant was removed and beads were resuspended in 300 µL 1x Polysome Buffer and a bead-polysome master mix was prepared of 500 µL per sample 1x Polysome Buffer and 10 µL washed beads. 510 µL of the bead-polysome buffer master mix was added to each ∼700 µL fraction with regular vortexing of the master mix. Samples were incubated on the tumbler at 4°C in the cold room at 15 rpm overnight. The next day, samples were centrifuged for 3 min at 1,000 rcf at 4°C. The sucrose pellets with the beads. 90% of the supernatant was removed with a pipette without touching the beads to not remove all sucrose with the first wash. 250 µL 1x Polysome Buffer was added for the first wash, samples were tumbled in the cold room for 3 min on the tumbler at 4°C, centrifuged for 3 min at 1,000 rcf at 4°C, and supernatant was discarded. A second wash with 250 µL 1x Polysome Buffer without tumbling but direct centrifugation was performed. All supernatant was removed and any residual liquid was removed from the beads with a small tip. 50 µL 2x SDS sample buffer (120 mM Tris-HCl pH 6.8, 4% SDS, 200 mM DTT, 20% glycerol, trace bromophenol blue, in nuclease-free water) was added to the bead pellets and samples were vortexed for 1-2 min for full resuspension. Samples were heated at 95°C for 5 min and equal volumes of 15 µL of 50 µL samples were loaded onto polyacrylamide SDS-PAGE gradient gels for Western blot analysis.

### Western blotting of protein from cultured cells

Lysates were prepared by resuspending cells in 1x PBS and pelleting by centrifugation at 500 g for 5 min at 4°C. The cell-pellets were taken up in M-PER™ Mammalian protein extraction reagent (Thermo Scientific), supplemented with 1x cOmplete protease inhibitor cocktail (Roche), briefly vortexed for 10 seconds and incubated on ice for 15 min. Lysates were cleared by centrifugation for 15 min and protein concentration of the supernatant was estimated by measuring absorption at 280 nm using a NanoPhotometer (Implen). Sample concentrations were normalized and diluted in 4x loading dye (1 M TRIS HCl pH 6.8, 350 mM SDS, 50% glycerol, 25% β-mercaptoethanol and trace-amounts of bromophenol blue). For electrophoresis, samples were loaded on a 12.5% SDS gel and run in 1x SDS running buffer (248 mM TRIS base, 1.92 M glycine, 1% SDS) at room temperature at 120 V (constant voltage) for 80 min. For western blotting, proteins were transferred to polyvinylidene difluoride (PVDF) membranes for one hour at 100 V and 4°C in transfer buffer (25 mM Tris, 192 mM glycine, 20% ethanol [v/v]). The membranes were subsequently blocked in 5% skimmed milk in TBST for 1 h, then incubated with the antibodies diluted in 5% skimmed milk in TBST overnight on a rocking shaker at 4°C. Membranes were washed three times in 1x TBST for 5 min each, incubated with the secondary antibody at room temperature for 1 h and washed 3 times in 1x TBST. Western blots were developed using Amersham™ ECL Prime Western Blotting Detection Reagent (Cytiva) at the Amersham imager 600 (Cytiva). For a list of commercial antibodies used, see Table S5.

In the Leppek lab, proteins were resolved on self-made 4-20% polyacrylamide gradient Tris-glycine SDS-PAGE gels and transferred onto 0.2 µm pore size PVDF membranes (Biorad) using the semi-dry Trans-Blot Turbo system (Biorad). Membranes were then blocked in 1x PBS-0.1% Tween-20 containing 5% non-fat milk powder for 1 h, incubated with primary antibodies diluted in the same solution for 1 h at RT or overnight at 4°C, and washed four times for 5 min in 1x PBS-0.1% Tween-20, incubated with secondary antibodies for 1 h in 1x PBS-0.1% Tween-20 and washed four times for 15 min in 1x PBS-0.1% Tween-20. Horseradish peroxidase-coupled secondary antibodies anti-rabbit (Amersham, Cytiva) in combination with Clarity Western ECL Substrate (Biorad) and imaging on an ImageQuant 800 (Amersham, Cytiva) were used for detection. Antibodies were diluted in 1x PBS-0.1% Tween-20 at 1:1,000-1:3,000 dilution in 5% BSA (w/v) in 1x PBS-A (azide).

### Statistics and reproducibility

All gel-based experiments were performed as three or four replicates. No statistical method was used to predetermine sample size. No data were excluded from the analyses. The experiments were not randomized, and the investigators were not blinded to allocation during experiments and outcome assessment. All datapoints represent measurements from distinct samples. Statistical significances (two-tailed P value) were determined by unpaired t-test, unless stated otherwise (defined significance levels: p<0.05 = *, p<0.01 = **, p<0.001 = ***, p<0.0001= ****), using Prism 10 software (GraphPad). Standard deviation (SD) is shown with error bars.

## ACKNOWLEDGEMENTS

We thank Klemens Wild and Astrid Hendricks for help with ribosome preparations, we thank Laura Hölzli for technical assistance; we thank Monika Langlotz of the FACS core facility at Heidelberg University for help with flow cytometry; we thank Janina L. Gerber, Sandra Köhler, Ege Çığırgan, Rebekka Arnold, Shikana Browne, Sabrina Simmet and Juliane Groß for their support. We thank Selene Cordeiro for technical support with buffers, gels, media and agar plates. We thank the Flow Cytometry Core Facility of the Medical Faculty at the University of Bonn, Germany, for providing help, services and devices funded by the Deutsche Forschungsgemeinschaft (DFG, German Research Foundation) - project number 216372545.

## Funding

This work was supported by the Deutsche Forschungsgemeinschaft (DFG, German Research Foundation): Emmy Noether Program, project number 442512666 and TRR319/A06, project number 439669440 (to J.P.), TRR319/B01 and C02, project number 439669440 (to C.D.), the Cluster of Excellence ImmunoSensation3 under Germany’s Excellence Strategy – EXC2151 – 390873048 (to K.L.); the Chica and Heinz Schaller Foundation (to J.P.); the Klaus Tschira Stiftung, grant #00.013.2021 (to C.D.); the Human Frontier Science Program (HFSP) Early Career Award, RGEC32/2023 (to K.L.); start-up funds of the Medical Faculty at the University Hospital Bonn and University of Bonn, Germany (to K.L.).

## Author contributions

A.N.W. performed almost all biochemical and all cell biological experiments. I.S.N.d.V. performed all nanopore sequencing experiments, I.S.N.d.V. and C.D. analyzed sequencing data. A.M.P. performed some biochemical experiments. K.L. performed and analyzed polysome profiling experiments. A.G. performed puromycin incorporation experiments. A.R. provided enzyme resources. C.D. and J.P. provided funding. J.P. conceived the study. The manuscript was written by A.N.W. and J.P. with contributions from all authors.

## Declaration of interests

The authors declare no competing interests.

## Materials and Correspondence

Correspondence and requests for materials should be addressed to

All sequencing data used in the analysis are available to any researcher for purposes of reproducing or extending the analysis under the following link: https://dataview.ncbi.nlm.nih.gov/object/PRJNA1446284?reviewer=7v8jjm8rgsf9k5tatpumpgl96i.

## Notes

https://dataview.ncbi.nlm.nih.gov/object/PRJNA1446284?reviewer=7v8jjm8rgsf9k5tatpumpgl96i

### Competing Interest Statement

The authors have declared no competing interest.

## REFERENCES

1. Shuman, S. (2023). RNA Repair: Hiding in Plain Sight. Annual Review of Genetics 57, 461–489. 10.1146/annurev-genet-071719-021856.

2. Cordes, J., Zhao, S., Engel, C.M., and Stingele, J. (2025). Cellular responses to RNA damage. Cell 188, 885–900. 10.1016/j.cell.2025.01.005.

3. Zaher, H.S., and Mosammaparast, N. (2025). RNA Damage Responses in Cellular Homeostasis, Genome Stability, and Disease. Annual Review of Pathology: Mechanisms of Disease 20, 433–457. 10.1146/annurev-pathmechdis-111523-023516.

4. Yan, L.L., and Zaher, H.S. (2019). How do cells cope with RNA damage and its consequences? Journal of Biological Chemistry 294, 15158–15171. 10.1074/jbc.REV119.006513.

5. Wurtmann, E.J., and Wolin, S.L. (2009). RNA under attack: Cellular handling of RNA damage. Critical Reviews in Biochemistry and Molecular Biology 44, 34–49. 10.1080/10409230802594043.

6. Simms, C.L., Hudson, B.H., Mosior, J.W., Rangwala, A.S., and Zaher, H.S. (2014). An Active Role for the Ribosome in Determining the Fate of Oxidized mRNA. Cell Reports 9, 1256–1264. 10.1016/j.celrep.2014.10.042.

7. Yan, L.L., Simms, C.L., McLoughlin, F., Vierstra, R.D., and Zaher, H.S. (2019). Oxidation and alkylation stresses activate ribosome-quality control. Nat Commun 10, 5611. 10.1038/s41467-019-13579-3.

8. Rahmanto, A.S., Blum, C.J., Scalera, C., Heidelberger, J.B., Mesitov, M., Horn-Ghetko, D., Gräf, J.F., Mikicic, I., Hobrecht, R., Orekhova, A., et al. (2023). K6-linked ubiquitylation marks formaldehyde-induced RNA-protein crosslinks for resolution. Molecular Cell 83, 4272–4289.e10.10.1016/j.molcel.2023.10.011.

9. Zhao, S., Cordes, J., Caban, K.M., Götz, M.J., Mackens-Kiani, T., Veltri, A.J., Sinha, N.K., Weickert, P., Kaya, S., Hewitt, G., et al. (2023). RNF14-dependent atypical ubiquitylation promotes translation-coupled resolution of RNA-protein crosslinks. Molecular Cell 83, 4290–4303.e9. 10.1016/j.molcel.2023.10.012.

10. LaRiviere, F.J., Cole, S.E., Ferullo, D.J., and Moore, M.J. (2006). A Late-Acting Quality Control Process for Mature Eukaryotic rRNAs. Molecular Cell 24, 619–626. 10.1016/j.molcel.2006.10.008.

11. Cole, S.E., LaRiviere, F.J., Merrikh, C.N., and Moore, M.J. (2009). A Convergence of rRNA and mRNA Quality Control Pathways Revealed by Mechanistic Analysis of Nonfunctional rRNA Decay. Molecular Cell 34, 440–450. 10.1016/j.molcel.2009.04.017.

12. Scully, R., Panday, A., Elango, R., and Willis, N.A. (2019). DNA double-strand break repair-pathway choice in somatic mammalian cells. Nat Rev Mol Cell Biol 20, 698–714. 10.1038/s41580-019-0152-0.

13. Li, Y., and Breaker, R.R. (1999). Kinetics of RNA Degradation by Specific Base Catalysis of Transesterification Involving the 2’-Hydroxyl Group. J. Am. Chem. Soc. 121, 5364–5372. 10.1021/ja990592p.

14. Breslow, R., and Huang, D.L. (1991). Effects of metal ions, including Mg2+ and lanthanides, on the cleavage of ribonucleotides and RNA model compounds. Proceedings of the National Academy of Sciences 88, 4080–4083. 10.1073/pnas.88.10.4080.

15. Jacobs, A.C., Resendiz, M.J.E., and Greenberg, M.M. (2010). Direct Strand Scission from a Nucleobase Radical in RNA. J. Am. Chem. Soc. 132, 3668–3669. 10.1021/ja100281x.

16. Paul, R., and Greenberg, M.M. (2016). Mechanistic Studies on RNA Strand Scission from a C2′-Radical. J. Org. Chem. 81, 9199–9205. 10.1021/acs.joc.6b01760.

17. Doma, M.K., and Parker, R. (2006). Endonucleolytic cleavage of eukaryotic mRNAs with stalls in translation elongation. Nature 440, 561–564. 10.1038/nature04530.

18. Guydosh, N.R., and Green, R. (2014). Dom34 Rescues Ribosomes in 3′ Untranslated Regions. Cell 156, 950–962. 10.1016/j.cell.2014.02.006.

19. Sinha, N.K., McKenney, C., Yeow, Z.Y., Li, J.J., Nam, K.H., Yaron-Barir, T.M., Johnson, J.L., Huntsman, E.M., Cantley, L.C., Ordureau, A., et al. (2024). The ribotoxic stress response drives UV-mediated cell death. Cell 187, 3652–3670.e40. 10.1016/j.cell.2024.05.018.

20. Iordanov, M.S., Pribnow, D., Magun, J.L., Dinh, T.-H., Pearson, J.A., Chen, S.L.-Y., and Magun, B.E. (1997). Ribotoxic Stress Response: Activation of the Stress-Activated Protein Kinase JNK1 by Inhibitors of the Peptidyl Transferase Reaction and by Sequence-Specific RNA Damage to the α-Sarcin/Ricin Loop in the 28S rRNA. Molecular and Cellular Biology 17, 3373–3381. 10.1128/MCB.17.6.3373.

21. Iordanov, M.S., Pribnow, D., Magun, J.L., Dinh, T.-H., Pearson, J.A., and Magun, B.E. (1998). Ultraviolet Radiation Triggers the Ribotoxic Stress Response in Mammalian Cells*. Journal of Biological Chemistry 273, 15794–15803. 10.1074/jbc.273.25.15794.

22. Greer, C.L., Peebles, C.L., Gegenheimer, P., and Abelson, J. (1983). Mechanism of action of a yeast RNA ligase in tRNA splicing. Cell 32, 537–546. 10.1016/0092-8674(83)90473-7.

23. Popow, J., Englert, M., Weitzer, S., Schleiffer, A., Mierzwa, B., Mechtler, K., Trowitzsch, S., Will, C.L., Lührmann, R., Söll, D., et al. (2011). HSPC117 Is the Essential Subunit of a Human tRNA Splicing Ligase Complex. Science 331, 760–764. 10.1126/science.1197847.

24. Gerber, J.L., Köhler, S., and Peschek, J. (2022). Eukaryotic tRNA splicing – one goal, two strategies, many players. Biological Chemistry 403, 765–778. 10.1515/hsz-2021-0402.

25. Popow, J., Schleiffer, A., and Martinez, J. (2012). Diversity and roles of (t)RNA ligases. Cell Mol Life Sci 69, 2657–2670. 10.1007/s00018-012-0944-2.

26. Chakravarty, A.K., Subbotin, R., Chait, B.T., and Shuman, S. (2012). RNA ligase RtcB splices 3′-phosphate and 5′-OH ends via covalent RtcB-(histidinyl)-GMP and polynucleotide-(3′)pp(5′)G intermediates. Proceedings of the National Academy of Sciences 109, 6072–6077. 10.1073/pnas.1201207109.

27. Desai, K.K., Bingman, C.A., Phillips, G.N., and Raines, R.T. (2013). Structures of the Noncanonical RNA Ligase RtcB Reveal the Mechanism of Histidine Guanylylation. Biochemistry 52, 2518–2525. 10.1021/bi4002375.

28. Popow, J., Jurkin, J., Schleiffer, A., and Martinez, J. (2014). Analysis of orthologous groups reveals archease and DDX1 as tRNA splicing factors. Nature 511, 104–107. 10.1038/nature13284.

29. Gerber, J.L., Morales Guzmán, S.I., Worf, L., Hubbe, P., Kopp, J., and Peschek, J. (2024). Structural and mechanistic insights into activation of the human RNA ligase RTCB by Archease. Nat Commun 15, 2378. 10.1038/s41467-024-46568-2.

30. Xu, Q., Teplow, D., Lee, T.D., and Abelson, J. (1990). Domain structure in yeast tRNA ligase. Biochemistry 29, 6132–6138.

31. Schwer, B., Sawaya, R., Ho, C.K., and Shuman, S. (2004). Portability and fidelity of RNA-repair systems. Proceedings of the National Academy of Sciences 101, 2788–2793. 10.1073/pnas.0305859101.

32. Sidrauski, C., Cox, J.S., and Walter, P. (1996). tRNA Ligase Is Required for Regulated mRNA Splicing in the Unfolded Protein Response. Cell 87, 405–413. 10.1016/S0092-8674(00)81361-6.

33. Lu, Y., Liang, F.-X., and Wang, X. (2014). A Synthetic Biology Approach Identifies the Mammalian UPR RNA Ligase RtcB. Molecular Cell 55, 758–770. 10.1016/j.molcel.2014.06.032.

34. Kosmaczewski, S.G., Edwards, T.J., Han, S.M., Eckwahl, M.J., Meyer, B.I., Peach, S., Hesselberth, J.R., Wolin, S.L., and Hammarlund, M. (2014). The RtcB RNA ligase is an essential component of the metazoan unfolded protein response. EMBO reports 15, 1278–1285. 10.15252/embr.201439531.

35. Jurkin, J., Henkel, T., Nielsen, A.F., Minnich, M., Popow, J., Kaufmann, T., Heindl, K., Hoffmann, T., Busslinger, M., and Martinez, J. (2014). The mammalian tRNA ligase complex mediates splicing of XBP1 mRNA and controls antibody secretion in plasma cells. The EMBO Journal 33, 2922–2936. 10.15252/embj.201490332.

36. Peschek, J., Acosta-Alvear, D., Mendez, A.S., and Walter, P. (2015). A conformational RNA zipper promotes intron ejection during non-conventional XBP1 mRNA splicing. EMBO reports 16, 1688–1698. 10.15252/embr.201540955.

37. Zhao, L.-W., Nardone, C., Chang, C., Paulo, J.A., Elledge, S.J., and Kennedy, S. (2025). An RNA splicing system that excises DNA transposons from animal mRNAs. Nature, 1–9. 10.1038/s41586-025-09853-8.

38. Nemudraia, A., Nemudryi, A., and Wiedenheft, B. (2024). Repair of CRISPR-guided RNA breaks enables site-specific RNA excision in human cells. Science 384, 808–814. 10.1126/science.adk5518.

39. Lindley, S.R., Subbaiah, K.C.V., Priyanka, F., Poosala, P., Ma, Y., Jalinous, L., West, J.A., Richardson, W.A., Thomas, T.N., and Anderson, D.M. (2024). Ribozyme-activated mRNA trans-ligation enables large gene delivery to treat muscular dystrophies. Science 386, 762–767. 10.1126/science.adp8179.

40. Engl, C., Schaefer, J., Kotta-Loizou, I., and Buck, M. (2016). Cellular and molecular phenotypes depending upon the RNA repair system RtcAB of Escherichia coli. Nucleic Acids Res 44, 9933–9941. 10.1093/nar/gkw628.

41. Temmel, H., Müller, C., Sauert, M., Vesper, O., Reiss, A., Popow, J., Martinez, J., and Moll, I. (2017). The RNA ligase RtcB reverses MazF-induced ribosome heterogeneity in Escherichia coli. Nucleic Acids Res 45, 4708–4721. 10.1093/nar/gkw1018.

42. Hughes, K.J., Chen, X., Burroughs, A.M., Aravind, L., and Wolin, S.L. (2020). An RNA Repair Operon Regulated by Damaged tRNAs. Cell Reports 33, 108527. 10.1016/j.celrep.2020.108527.

43. Tian, Y., Zeng, F., Raybarman, A., Fatma, S., Carruthers, A., Li, Q., and Huang, R.H. (2022). Sequential rescue and repair of stalled and damaged ribosome by bacterial PrfH and RtcB. Proceedings of the National Academy of Sciences 119, e2202464119. 10.1073/pnas.2202464119.

44. Hindley, H.J., Gong, Z., Moradian, S., Giuliano, M.G., Sapelkin, A., Kotta-Loizou, I., Buck, M., Engl, C., and Weiße, A.Y. (2025). Heterogeneity in responses to ribosome-targeting antibiotics mediated by bacterial RNA repair. Nat Commun 16, 9620. 10.1038/s41467-025-64759-3.

45. Nayak, S.K., Bagga, S., Gaur, D., Nair, D.T., Salunke, D.M., and Batra, J.K. (2001). Mechanism of Specific Target Recognition and RNA Hydrolysis by Ribonucleolytic Toxin Restrictocin. Biochemistry 40, 9115–9124. 10.1021/bi010923m.

46. Schindler, D.G., and Davies, J.E. (1977). Specific cleavage of ribosomal RNA caused by alpha sarcin. Nucleic Acids Res 4, 1097–1110.10.1093/nar/4.4.1097.

47. Endo, Y., and Wool, I.G. (1982). The site of action of alpha-sarcin on eukaryotic ribosomes. The sequence at the alpha-sarcin cleavage site in 28 S ribosomal ribonucleic acid. Journal of Biological Chemistry 257, 9054–9060. 10.1016/S0021-9258(18)34241-8.

48. Korennykh, A.V., Plantinga, M.J., Correll, C.C., and Piccirilli, J.A. (2007). Linkage between Substrate Recognition and Catalysis during Cleavage of Sarcin/Ricin Loop RNA by Restrictocin. Biochemistry 46, 12744–12756. 10.1021/bi700931y.

49. Banerjee, A., Ghosh, S., Goldgur, Y., and Shuman, S. (2019). Structure and two-metal mechanism of fungal tRNA ligase. Nucleic Acids Res 47, 1428–1439. 10.1093/nar/gky1275.

50. Peschek, J., and Walter, P. (2019). tRNA ligase structure reveals kinetic competition between non-conventional mRNA splicing and mRNA decay. eLife 8, e44199. 10.7554/eLife.44199.

51. Köhler, S., Kopp, J., Maiti, S., Bujnicki, J.M., and Peschek, J. (2025). Structure of fungal tRNA ligase Trl1 with RNA reveals conserved substrate-binding principles. Nat Struct Mol Biol 32, 1657–1668. 10.1038/s41594-025-01589-3.

52. Correll, C.C., Munishkin, A., Chan, Y.-L., Ren, Z., Wool, I.G., and Steitz, T.A. (1998). Crystal structure of the ribosomal RNA domain essential for binding elongation factors. Proceedings of the National Academy of Sciences 95, 13436–13441. 10.1073/pnas.95.23.13436.

53. Doris, S.M., Smith, D.R., Beamesderfer, J.N., Raphael, B.J., Nathanson, J.A., and Gerbi, S.A. (2015). Universal and domain-specific sequences in 23S–28S ribosomal RNA identified by computational phylogenetics. RNA 21, 1719–1730. 10.1261/rna.051144.115.

54. Szewczak, A.A., Moore, P.B., Chang, Y.L., and Wool, I.G. (1993). The conformation of the sarcin/ricin loop from 28S ribosomal RNA. Proceedings of the National Academy of Sciences 90, 9581–9585. 10.1073/pnas.90.20.9581.

55. Yuan, Y., Stumpf, F.M., Schlor, L.A., Schmidt, O.P., Saumer, P., Huber, L.B., Frese, M., Höllmüller, E., Scheffner, M., Stengel, F., et al. (2023). Chemoproteomic discovery of a human RNA ligase. Nat Commun 14, 842. 10.1038/s41467-023-36451-x.

56. Wassarman, D.A., and Steitz, J.A. (1991). Structural Analyses of the 7SK Ribonucleoprotein (RNP), the Most Abundant Human Small RNP of Unknown Function. Molecular and Cellular Biology 11, 3432–3445. 10.1128/mcb.11.7.3432-3445.1991.

57. Nguyen, V.T., Kiss, T., Michels, A.A., and Bensaude, O. (2001). 7SK small nuclear RNA binds to and inhibits the activity of CDK9/cyclin T complexes. Nature 414, 322–325. 10.1038/35104581.

58. Stark, B.C., Kole, R., Bowman, E.J., and Altman, S. (1978). Ribonuclease P: an enzyme with an essential RNA component. Proceedings of the National Academy of Sciences 75, 3717–3721. 10.1073/pnas.75.8.3717.

59. Simsek, D., Tiu, G.C., Flynn, R.A., Byeon, G.W., Leppek, K., Xu, A.F., Chang, H.Y., and Barna, M. (2017). The Mammalian Ribo-interactome Reveals Ribosome Functional Diversity and Heterogeneity. Cell 169, 1051–1065.e18. 10.1016/j.cell.2017.05.022.

60. Colognori, D.A., Wasko, K.M., Trinidad, M.I., Zhou, Z., and Doudna, J.A. (2026). Spligation enables programmable chimeric RNA generation in living cells. bioRxiv, 2026.03.06.709984. 10.64898/2026.03.06.709984.

61. Cherry, P.D., Peach, S.E., and Hesselberth, J.R. (2019). Multiple decay events target HAC1 mRNA during splicing to regulate the unfolded protein response. eLife 8, e42262. 10.7554/eLife.42262.

62. Peebles, C.L., Ogden, R.C., Knapp, G., and Abelson, J. (1979). Splicing of yeast tRNA precursors: a two-stage reaction. Cell 18, 27–35. 10.1016/0092-8674(79)90350-7.

63. Dremel, S.E., Koparde, V.N., Arbuckle, J.H., Hogan, C.H., Kristie, T.M., Krug, L.T., Conrad, N.K., and Ziegelbauer, J.M. (2025). Noncanonical circRNA biogenesis driven by alpha and gamma herpesviruses. EMBO J 44, 2323–2352. 10.1038/s44318-025-00398-0.

64. Khatter, H., Myasnikov, A.G., Natchiar, S.K., and Klaholz, B.P. (2015). Structure of the human 80S ribosome. Nature 520, 640–645. 10.1038/nature14427.

65. Khatter, H., Myasnikov, A.G., Mastio, L., Billas, I.M.L., Birck, C., Stella, S., and Klaholz, B.P. (2014). Purification, characterization and crystallization of the human 80S ribosome. Nucleic Acids Research 42, e49–e49. 10.1093/nar/gkt1404.

66. Matsuura-Suzuki, E., Toh, H., and Iwasaki, S. (2023). Human-rabbit Hybrid Translation System to Explore the Function of Modified Ribosomes. BIO-PROTOCOL 13. 10.21769/BioProtoc.4714.

67. Van Der Toorn, W., Bohn, P., Liu-Wei, W., Olguin-Nava, M., Gribling-Burrer, A.-S., Smyth, R.P., and Von Kleist, M. (2025). Demultiplexing and barcode-specific adaptive sampling for nanopore direct RNA sequencing. Nat Commun 16, 3742. 10.1038/s41467-025-59102-9.

68. Leger, A., Amaral, P.P., Pandolfini, L., Capitanchik, C., Capraro, F., Miano, V., Migliori, V., Toolan-Kerr, P., Sideri, T., Enright, A.J., et al. (2021). RNA modifications detection by comparative Nanopore direct RNA sequencing. Nat Commun 12, 7198. 10.1038/s41467-021-27393-3.

69. Naarmann-de Vries, I.S., Zorbas, C., Lemsara, A., Piechotta, M., Ernst, F.G.M., Wacheul, L., Lafontaine, D.L.J., and Dieterich, C. (2023). Comprehensive identification of diverse ribosomal RNA modifications by targeted nanopore direct RNA sequencing and JACUSA2. RNA Biology 20, 652–665. 10.1080/15476286.2023.2248752.

70. Argüello, R.J., Combes, A.J., Char, R., Gigan, J.-P., Baaziz, A.I., Bousiquot, E., Camosseto, V., Samad, B., Tsui, J., Yan, P., et al. (2020). SCENITH: A Flow Cytometry-Based Method to Functionally Profile Energy Metabolism with Single-Cell Resolution. Cell Metabolism 32, 1063–1075.e7. 10.1016/j.cmet.2020.11.007.

