## Supplemental material for "Eukaryotic tRNA ligases mediate RNA break repair"

**The PDF file includes:**

Fig. S1-S5

Tables S1-S5

**Other Supplementary Materials for this manuscript include the following:**

Movie S1

Source data

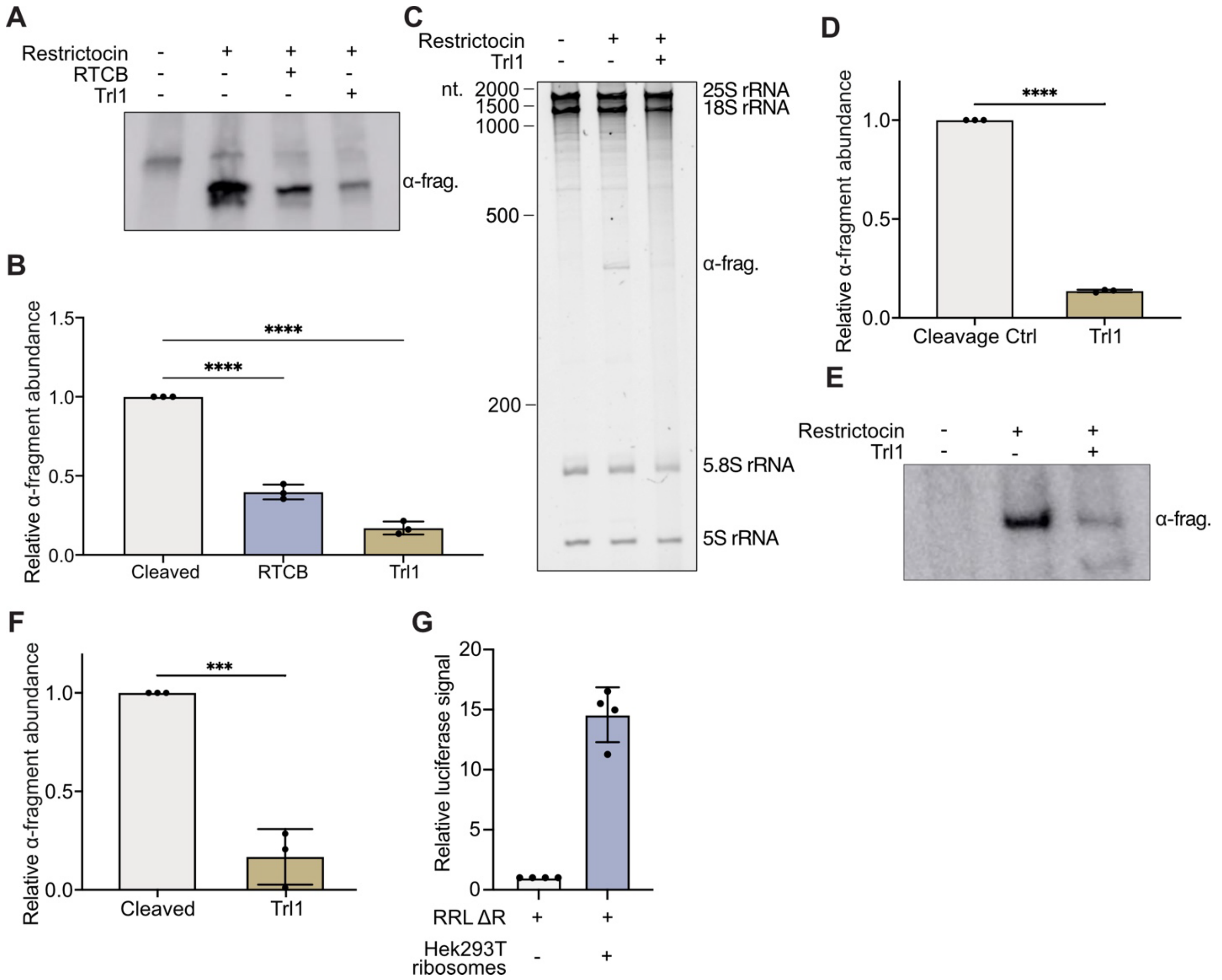


**Figure S1**

(A) Northern blot against α-fragment using samples from Figure 2C.
(B) Quantification of S1A. n=3 independent replicates.
(C) Urea-Page showing rRNA isolated from 80S yeast ribosomes before, after cleavage with restrictocin and after repair with fungal Trl1 ligase.
(D) Quantification of S1C. n=3 independent replicates.
(E) Northern blot against α-fragment with samples from S1C.
(F) Quantification of S1E. n=3 independent replicates.
(G) In vitro translation assay using ribosome depleted rabbit reticulocyte lysate (RRL) with and without unmodified HEK293T ribosomes. Luciferase signal for samples supplemented with HEK293T ribosomes is normalized to the sample without HEK293T ribosomes. n=4 independent replicates. Statistical significances for all panels in Figure S1 were determined using unpaired t-test (defined significance levels: $p<0.05$ = *, $p<0.01$ = **, $p<0.001$ = ***, $p<0.0001$= ****). Error bars depict the standard deviation (SD).

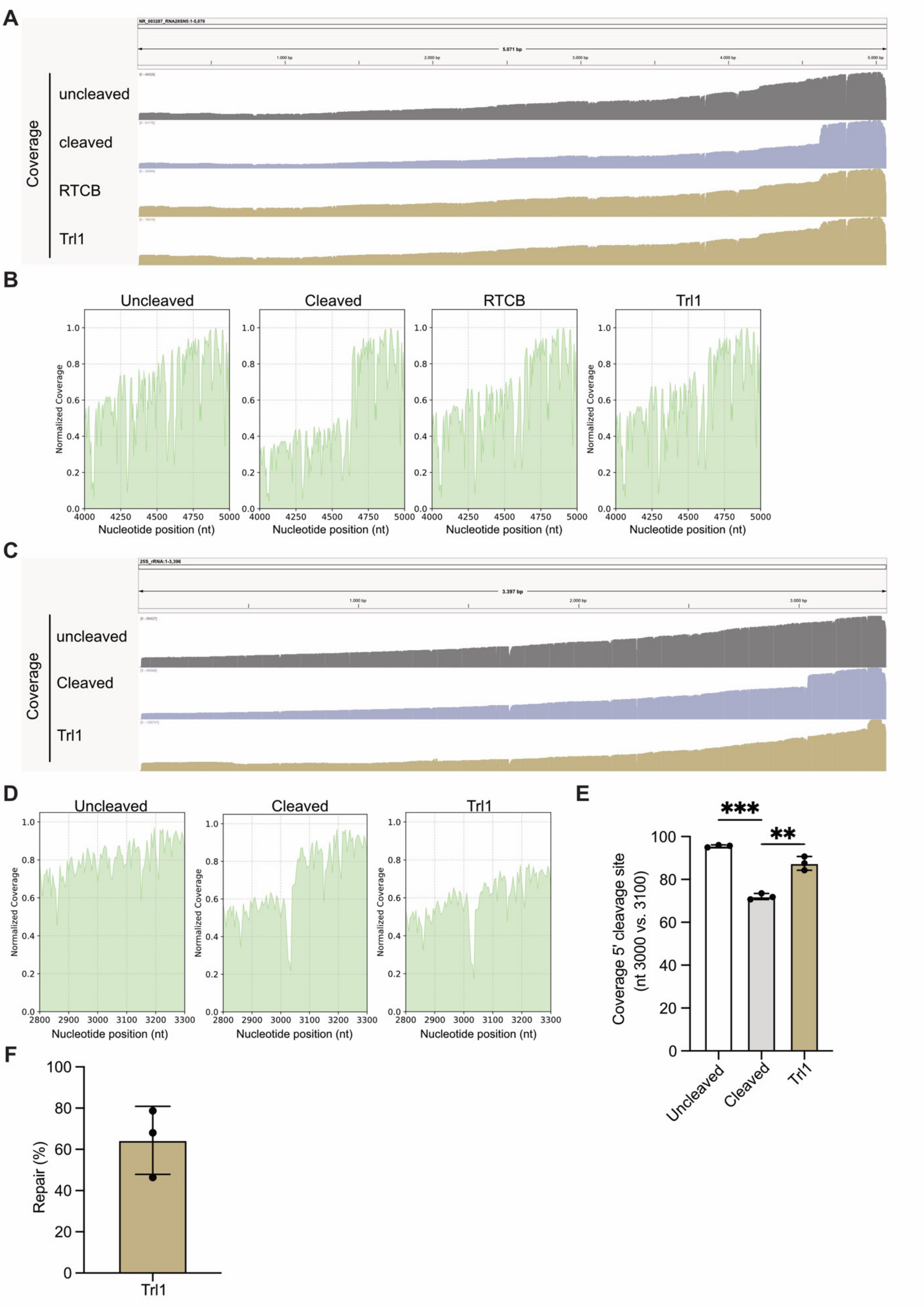
A
NR_003287_RNA28SN5:1-5,070
5,071 bp
1,000 bp
2,000 bp
3,000 bp
4,000 bp
5,000 bp
Coverage
uncleaved
cleaved
RTCB
Trl1
B
Uncleaved
Cleaved
RTCB
Trl1
Normalized Coverage
Nucleotide position (nt)
C
25S_rRNA:1-3,396
3,397 bp
1,000 bp
2,000 bp
3,000 bp
Coverage
uncleaved
Cleaved
Trl1
D
Uncleaved
Cleaved
Trl1
Normalized Coverage
Nucleotide position (nt)
E
***
**
Coverage 5' cleavage site
(nt 3000 vs. 3100)
Uncleaved
Cleaved
Trl1
F
Repair (%)
Trl1

**Figure S2**

(A) IGV sequence coverage plots of rRNA sequenced with nanopore sequencing, corresponding to Figure 3A. Intact HEK293T ribosomes were either left untreated, or specifically cleaved with restrictocin and repaired by either RTCB or Trl1. The drop in sequence coverage at around 4500 nt is strongest in the cleaved samples and reduced in both the RTCB- and Trl1-repaired samples.
(B) Zoom ins to the region of the restrictocin cleavage site (position 4000-5000). Coverage was normalized to the highest coverage peaks.
(C) IGV sequence coverage plots of yeast rRNA sequenced with nanopore sequencing. Intact yeast ribosomes were either left untreated, or specifically cleaved with restrictocin and repaired by Trl1.
(D) Zoom ins to the region of the restrictocin cleavage site (2800 nt – 3300 nt). Coverage was normalized to the highest coverage peaks. The drop in sequence coverage at around 3000nt is strongest in the cleaved samples and reduced in both the RTCB and Trl1 repaired samples. An additional drop in sequence coverage at around 3400 nt is present in all samples, but most prominent in the Trl1 ligated sample.
(E) Quantification of the sequence coverage drop by normalizing the sequence coverage 5′ of the cleavage site (nt 3000) to the sequence coverage 3′ of the cleavage site (nt 3100) for all analyzed samples. n=3 independent replicates. Statistical significance of cleavage and repair was determined by One-way ANOVA followed by Tukey's multiple comparison test (defined significance levels: $p<0.05$ = *, $p<0.01$ = **, $p<0.001$ = ***)
(F) Calculation of the percentage of cleaved HEK293T ribosomes that are successfully repaired by Trl1. Error bars depict the standard deviation (SD).

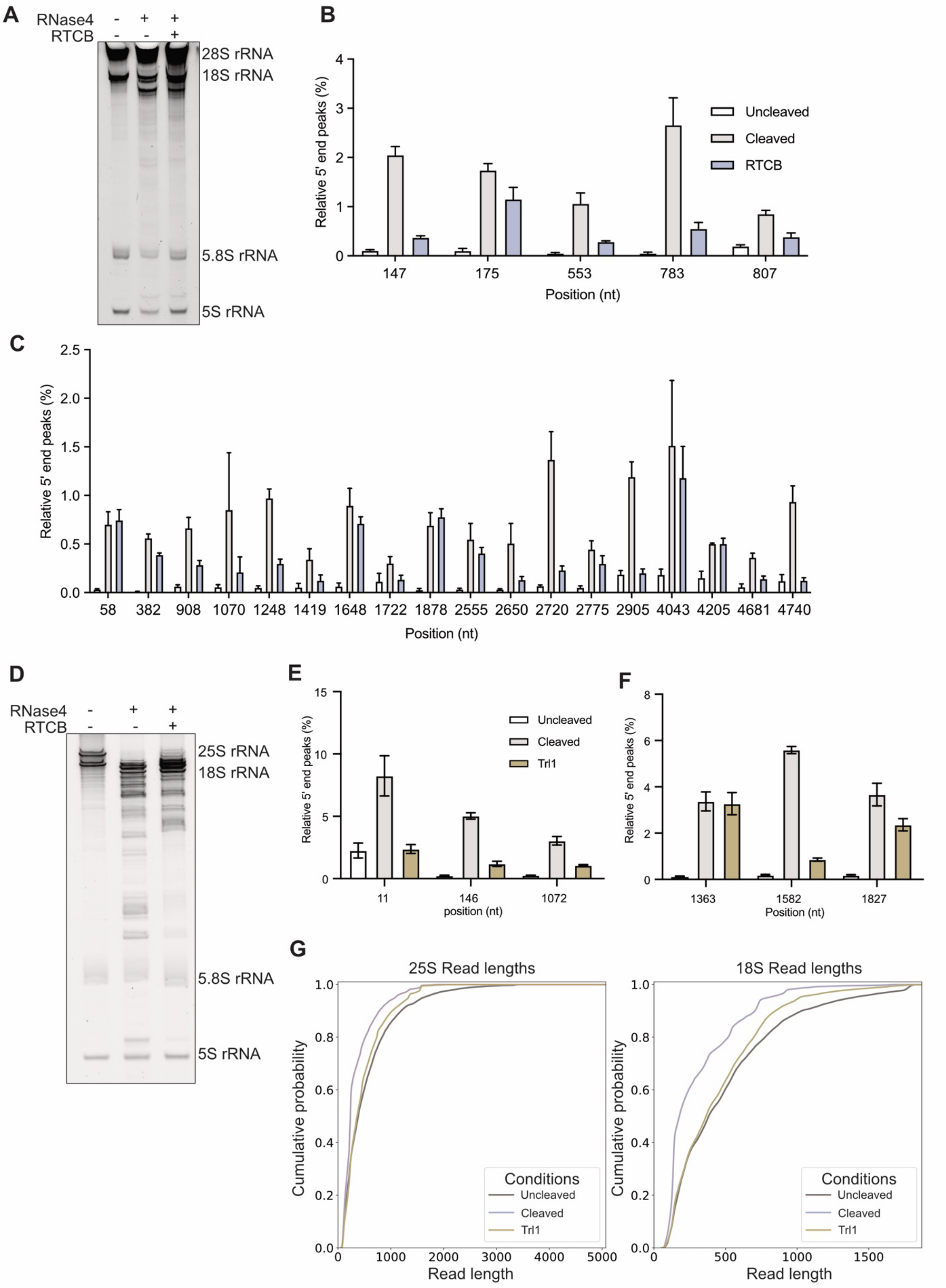
A
RNase4 - + +
RTCB - - +
28S rRNA
18S rRNA
5.8S rRNA
5S rRNA
B
Relative 5' end peaks (%)
Uncleaved
Cleaved
RTCB
147
175
553
783
807
Position (nt)
C
Relative 5' end peaks (%)
58 382 908 1070 1248 1419 1648 1722 1878 2555 2650 2720 2775 2905 4043 4205 4681 4740
Position (nt)
D
RNase4 - + +
RTCB - - +
25S rRNA
18S rRNA
5.8S rRNA
5S rRNA
E
Relative 5' end peaks (%)
Uncleaved
Cleaved
Trl1
11
146
1072
position (nt)
F
Relative 5' end peaks (%)
1363
1582
1827
Position (nt)
G
25S Read lengths
18S Read lengths
Cumulative probability
Read length
Conditions
Uncleaved
Cleaved
Trl1

**Figure S3**

(A) Urea-PAGE showing rRNA isolated from 80S HEK293T ribosomes, before and after RNase4 treatment, as well as after RNA break repair with RTCB.
(B and C) Relative 5′-end peaks in 18S and 28S rRNA detected by nanopore sequencing of the samples shown in S3A n=3 independent replicates.
(D) Urea-PAGE showing rRNA isolated from 80S yeast ribosomes, before and after RNase4 treatment, as well as after RNA break repair with Trl1.
(E and F) Relative 5′-end peaks in yeast 18S and 25S rRNA detected by nanopore sequencing of the samples shown in S3D. n=3 independent experiments.
(G) eCDF plots of cumulative probability for each read length after nanopore sequencing of rRNA from ribosomes untreated yeast ribosomes, yeast ribosomes treated with RNase4 and yeast ribosomes repaired with Trl1. Shown are read lengths for 25S rRNA and 18S. Error bars for all panels in this figure depict the standard deviation (SD).

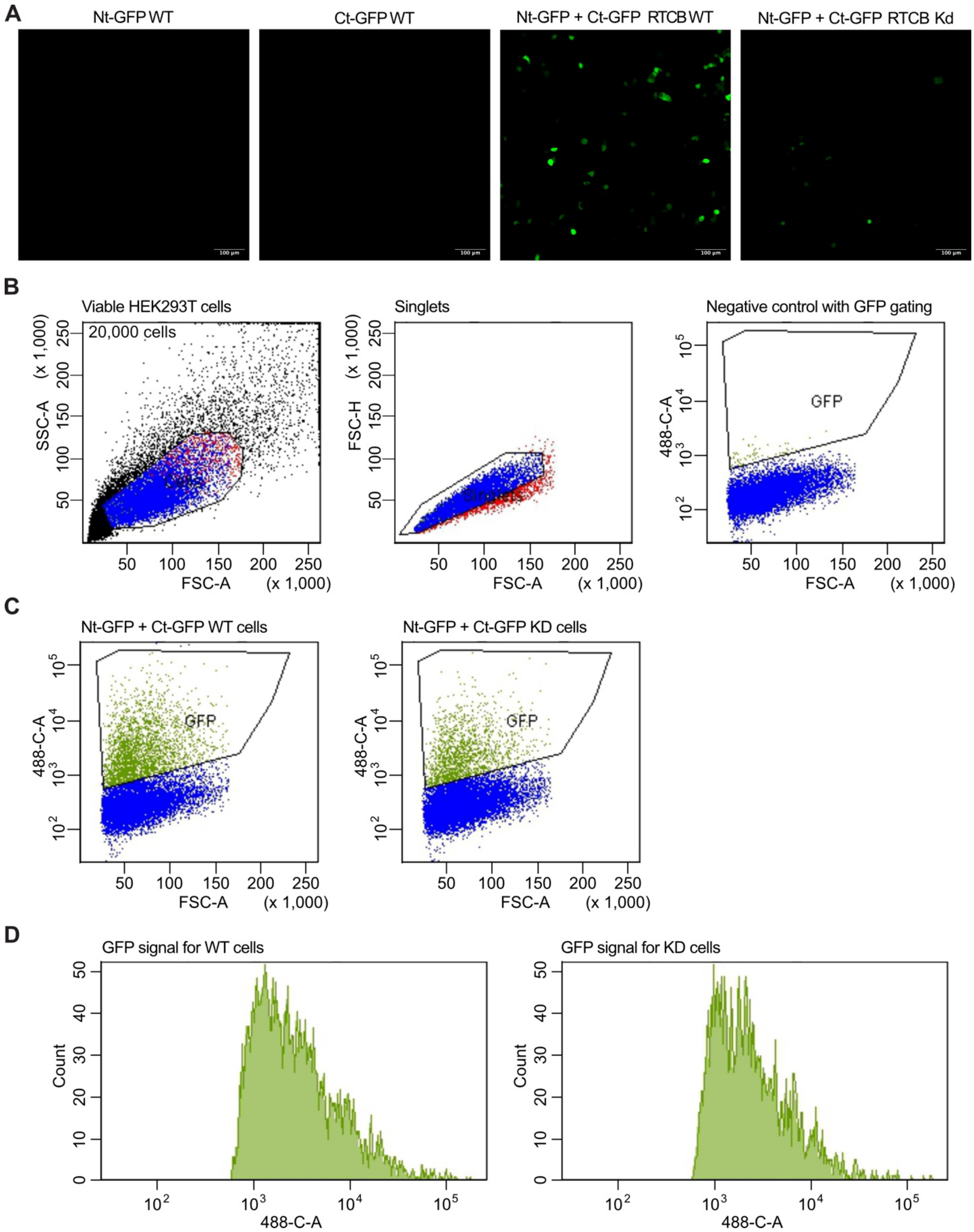


**Figure S4**

(A) Left: Widefield microscopy images (20x) of HEK293T WT cells either transfected with only the Nt-GFP-Twister plasmid, the Rbz-HH-Ct-GFP plasmid, or co-transfected with both

plasmids. Right: Widefield microscopy images of HEK293T RTCB KD cells co-transfected with both plasmids. All images were taken with the same microscope settings and were adjusted with the same settings. Scalebar set to 100 µm.
(B) Gating strategy for flow cytometry experiments. With the first gate, only viable HEK293T were selected, with the second gate, only singlets were selected. The third gate was chosen so that only GFP-positive cells would be selected, shown here is a representative image of the negative control, transfected with only the Nt-GFP half.
(C) Flow cytometry panels showing the results for a representative replicate for both the RTCB WT and RTCB KD cells co-transfected with Nt-GFP and Ct-GFP.
(D) Analysis of GFP (488 nm) signal for a representative replicate for both the RTCB WT and RTCB KD cells co-transfected with Nt-GFP and Ct-GFP.

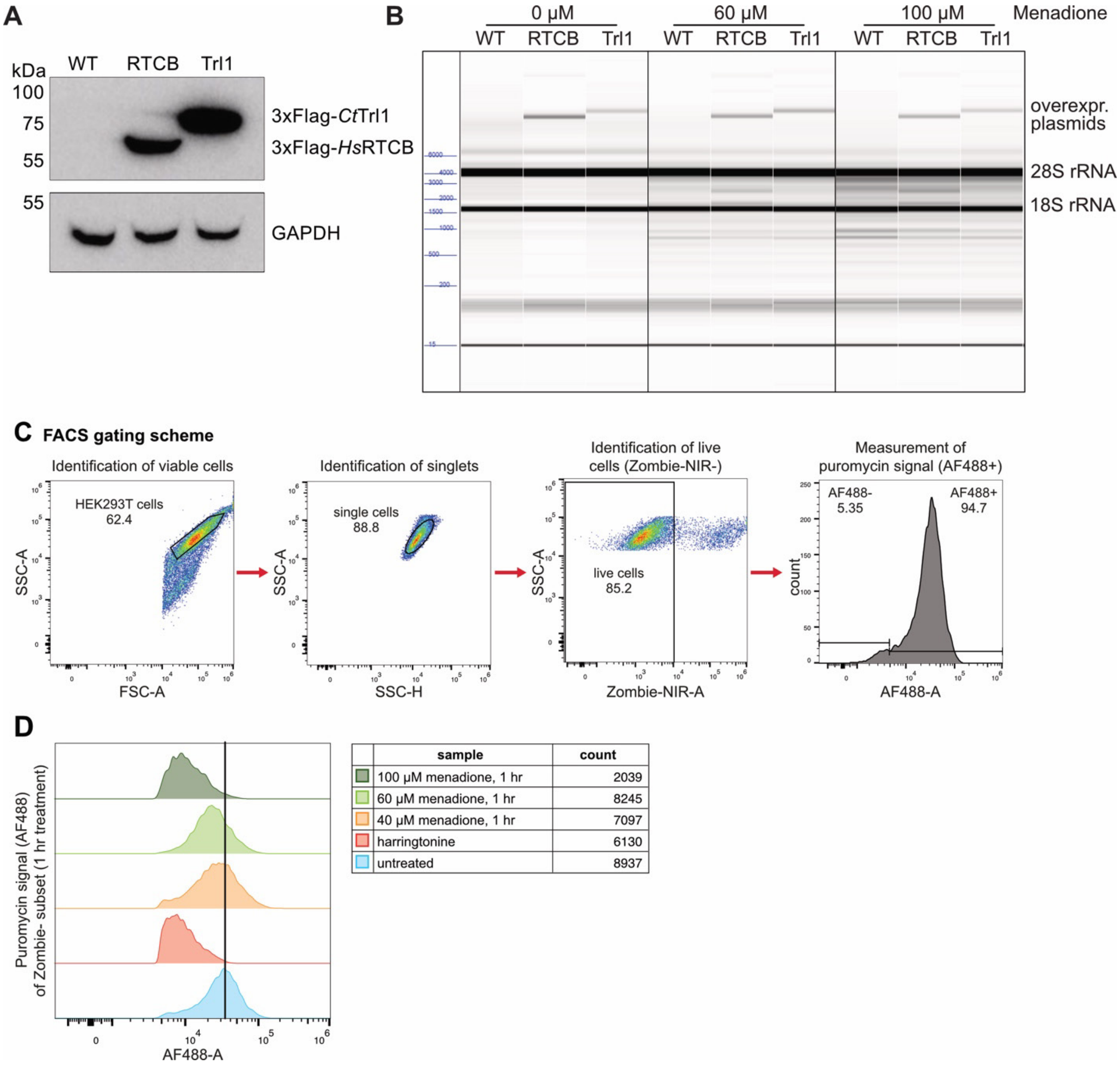


**Figure S5**

(A) Western blot using anti-Flag antibody with cell lysate from HEK293T WT cells, and cells with either 3xFlag-*hs*RTCB, or 3xFlag-*ct*Trl1 overexpression plasmids.
(B) Electropherograms showing the effect of the RTCB and Trl1 overexpression on the cellular RNA integrity with increasing concentrations of menadione (0 µM, 60 µM, 100 µM), (n=2).
(C) Gating strategy for flow cytometry analysis of the puromycin incorporation assay.
(D) Results from flow cytometry analysis for the puromycin incorporation assay. Left: AF488-A is plotted against the puromycin signal. Right: The cell count for each condition is listed.

**Table S1.** Plasmids used in this study.

| Name | Resistance | Source |
|---|---|---|
| pET24A ompA-Restrictocin-His6 | Kan | This study |
| pcDNA3.1(+)-3xFlag-*Hs*RTCB | Amp/Neo | This study |
| pcDNA3.1(+)-3xFlag-*Ct*Trl1 | Amp/Neo | This study |
| pcDNA3.1(+)-NtGFP-Twister | Amp/Neo | This study |
| pcDNA3.1(+)-HH-CtGFP | Amp/Neo | This study |
| pET15b-His6-ctTrl1 | Amp | Peschek and Walter 2019 [1] |
| pFastBac HT B-*Hs*RTCB | Amp/Genta | Gerber et al. 2024[2] |
| pET28a-*Hs*Archease | Kan | Gerber et al. 2024[2] |

**Table S2.** Restriction enzyme cloning primers used in the study.

| Name | Sequence | Source |
|---|---|---|
| pcDNA3.1(+)-3xFLAG-*Hs*RTCB cloning fwd | GTACgaattcAGATTACGATATCCCAACGACCATGAGTCGCAGCTATAATGATG | Merck/Sigma |
| pcDNA3.1(+)-3xFLAG-*Hs*RTCB cloning rev | GTACaagcttCTATCCTTTGATCACAGCAATTGGTCTCAG | Merck/Sigma |
| pcDNA3.1(+)-3xFLAG-*Ct*Trl1 cloning fwd | CTAGAATTCAGATTACGATATCCCAACGACCATGGATGGTACCGCCGAGAAGC | Merck/Sigma |
| pcDNA3.1(+)-3xFLAG-*Ct*Trl1 cloning rev | CTAGAAGCTTCTACCTTGACAACACACCCTTGACGATGCCCC | Merck/Sigma |

**Table S3.** Digoxigenin-labelled Northern probes used in this study.

| Name | Sequence | Source |
|---|---|---|
| Human α-fragment probe | GACTGCTCTGCTACGTACGA(DIG) | Merck/Sigma |
| Yeast α-fragment probe | ACAAATCAGACAACAAAGGCTTAA(DIG) | Merck/Sigma |

**Table S4.** RNA oligonucleotide used in this study.

| Name | Sequence | Source |
|---|---|---|
| SRL-oligonucleotide | UUGCGCUCCUCAGUACGAGAGGAACCGGAGCGC | Merck/Sigma |

**Table S5.** commercial antibodies used in this study.

| Antibody name | Species | dilution | Company | Catalog |
|---|---|---|---|---|
| DYKDDDDK, anti-FLAG | mouse | 1:10,000 | Proteintech | 66008-4-Ig |
| RtcB Specific antibody | rabbit | 1:3000 | Proteintech | 19809-1-AP |
| HRP-conjugated GAPDH | rabbit | 1:10,000 | Proteintech | HRP-60004 |
| Anti-mouse IgG HRP Conjugate | goat | 1:3000 | Promega | W4028 |
| Anti rabbit HRP | goat | 1:10,000 | Proteintech | SA00001-2 |
| rabbit anti-RPS6/eS6 | rabbit | 1:3000 | Cell Signaling | 5G10/2217L |
| Rabbit anti-RPL10A/uL1 | Rabbit | 1:3000 | Abcam | ab174318 |
| anti-Puro, clone 12D10, Alexa Fluor 488 | mouse | 1:1000 | Merck/Sigma | MABE343-AF488 |
| anti-Puromycin, clone 12D10, Alexa Flour 647 | mouse | 1:1000 | Merck/Sigma | MABE343-AF647 |

**Movie S1.**

Structural movie of the human ribosome (PDB 4ug0)[3]. Shown in dark violet is the SRL as a reference point, the breakage sites identified by either analysis are shown in dark red, the space in between the mapped break sites is shown in light green.

**Supplemental references:**


1. Peschek, J., and Walter, P. (2019). tRNA ligase structure reveals kinetic competition between non-conventional mRNA splicing and mRNA decay. eLife *8*, e44199. https://doi.org/10.7554/eLife.44199.

2. Gerber, J.L., Morales Guzmán, S.I., Worf, L., Hubbe, P., Kopp, J., and Peschek, J. (2024). Structural and mechanistic insights into activation of the human RNA ligase RTCB by Archease. Nat. Commun. *15*, 2378. https://doi.org/10.1038/s41467-024-46568-2.

3. Khatter, H., Myasnikov, A.G., Natchiar, S.K., and Klaholz, B.P. (2015). Structure of the human 80S ribosome. Nature *520*, 640–645. https://doi.org/10.1038/nature14427.
